# Smaller Hippocampal Volume among Black and Latinx Youth Living in High-Stigma Contexts

**DOI:** 10.1101/2020.10.09.333328

**Authors:** Mark L Hatzenbuehler, David G Weissman, Sarah McKetta, Micah R Lattanner, Jessie V Ford, Deanna M Barch, Katie A McLaughlin

**Affiliations:** Harvard University; Columbia University; Washington University

**Keywords:** stigma, hippocampal volume, neurodevelopment, context effects

## Abstract

**Background:** To determine whether being raised in a stigmatizing context influences neurodevelopment in children.

**Methods:** We drew data from one of the first national, multi-site neuroimaging studies with substantial variability in sociopolitical contexts: the Adolescent Brain and Cognitive Development study (N=11,534; *M*=9.9 years). We measured structural stigma surrounding gender, race, and Latinx ethnicity using objective state-level indicators of social policies and prejudicial attitudes and individual-level experiences of perceived discrimination, and examined associations with two neural markers: hippocampal volume and amygdala reactivity to threat.

**Results:** In a pre-registered analysis, we find that Black (*B*=−57.27, *p*=0.025) and Latinx (*B*=−41.02, *p*=0.037) youths raised in higher-stigma environments have smaller hippocampal volume than youths raised in low-stigma contexts, controlling for demographics and family socioeconomic status. This association was marginal among girls (*p*=0.082). The magnitude of the observed difference in hippocampal volume between high and low structural stigma contexts was equivalent to a $20,000 difference in annual family income. Stigmatizing environments were unrelated to hippocampal volume in non-stigmatized youth, providing evidence of specificity. Perceived discrimination was unrelated to hippocampal volume among any of the stigmatized groups. No associations between individual or structural forms of stigma and amygdala reactivity to threat were observed in the stigmatized groups.

**Conclusions:** We provide novel evidence that an objective measure of stigma at the contextual level (structural stigma) may have a stronger influence on neurodevelopment than subjective perceptions of stigma measured at the individual level, suggesting that contextual approaches may yield new insights into neurodevelopment.

## Introduction

Stigma—defined as the co-occurrence of labeling, stereotyping, status loss, and discrimination in a context in which power is exercised (1)—contributes to adverse mental health outcomes for marginalized groups through its influence on processes across individual, interpersonal, and structural levels (1–3). At the individual level, stigma manifests as psychological responses through which stigmatized individuals perceive and react to stigma, including identity concealment (4), self-stigma (5), and expectations of rejection (6). Interpersonal forms of stigma refer to interactional processes that occur between the stigmatized and non-stigmatized, such as discriminatory treatment (7). Although most research has focused on the mental health consequences of stigma at the individual and interpersonal levels (7), growing evidence indicates that structural stigma—defined as “societal-level conditions, cultural norms and institutional policies and practices that constrain the opportunities, resources, and wellbeing of the stigmatized”(8)—represents an additional risk factor for psychopathology among the stigmatized (3). For instance, observational studies have shown that individuals living in states with fewer (vs. greater) legal protections for their stigmatized group (e.g., restrictive immigration policies) have higher levels of psychological distress (9,10) and psychiatric disorders (11). Further, quasi-experimental studies have demonstrated that rates of mental disorders (12) and psychological distress (13) increase among stigmatized populations following increases in structural stigma (e.g., passage of laws denying services to same-sex couples).

Despite consistent evidence for the adverse mental health consequences of stigma, the biological mechanisms through which stigma contributes to risk for psychopathology are only beginning to be understood (3,14). Experimental and observational studies have documented a variety of physiological responses to stigma-related experiences, including changes in immune functioning, inflammatory processes, and regulation of the hypothalamic-pituitary-adrenal (HPA) axis (15,16). Surprisingly, although social factors like rejection, exclusion, and early-life adversity (e.g., childhood trauma) have been associated consistently with brain structure and function (17–30), few studies have examined neurodevelopmental sequelae of stigma.

The current study begins to address this gap in the literature. Specifically, we examine whether individual and structural forms of stigma are associated with two neural markers: hippocampal volume and amygdala reactivity to threat. We chose these outcomes because they are associated with stress exposure (19,25–31), consistent with social identity threat theories of stigma that conceptualize it as a chronic stressor (7). Additionally, both neural markers are associated with multiple forms of psychopathology (32–38), and thus may serve as mechanisms linking stigma with mental health problems (7,8). To address our research question, we required a unique data structure that not only included measures of stigma at the individual level, but also sampled respondents from a range of social environments that differed in structural stigma. This presented a methodological challenge, because most neuroimaging studies are conducted in one (or a small number) of communities. In such designs, respondents are similarly exposed to the same macro-social context—known as a ubiquitous exposure (39)—precluding the possibility of linking contextual variation with neural outcomes. Fortunately, neuroimaging studies with meaningful variation in social context have recently become available. In this study, we use data from one of the first national, multi-site neuroimaging studies with substantial variability in sociopolitical contexts: the Adolescent Brain and Cognitive Development (ABCD) study. This dataset measured brain structure and function in 11,534 youth sampled from 17 states, affording significant geographic variability in exposure to stigmatizing contexts among youth.

We examined three stigmatized groups—girls, racial (Black), and ethnic (Latinx) minority youth (mean age=9.9 years)—informed by the developmental literature on identity awareness, formation, and responsivity to identity-based stressors, which indicates that gender (40), racial (41), and ethnic (42) identities emerge during early childhood. Girls, Black, and Latinx youth report identity awareness and constancy, and perceptions of group-based stereotypes, by 9-10 years, the age of the baseline ABCD sample. We tested two pre-registered hypotheses. First, we predicted that greater exposure to individual and structural stigma would be associated with smaller hippocampal volume and elevated amygdala reactivity to threat cues (i.e., fearful relative to neutral faces) among girls, Black, and Latinx youth, controlling for demographics and family socioeconomic status (SES). Second, we hypothesized that structural stigma would be unrelated to hippocampal volume or amygdala reactivity in the non-stigmatized comparison groups: boys, Whites, and non-Hispanic Whites. This analysis serves as a negative control approach (43), in that we test whether there is an association among the groups where we would not theoretically expect it. Because structural stigma represents a distal social factor, we expected that the effect sizes for this variable would be small in magnitude as compared to proximal social factors (e.g., childhood trauma) that have been the focus of most neuroimaging research (17–22). Nonetheless, even small effect sizes can be substantively meaningful (44), especially when they relate to factors whose potential influence is scaled over large populations (45).

## Methods

### Sample

Data come from the Adolescent Brain and Cognitive Development (ABCD) study, the largest study of brain development in the U.S. (https://abcdstudy.org). We drew data from the Year 1 assessment (ABCD 2.0) of 11,534 youth. Twenty-one study sites were included from across the U.S. From these sites, a stratified probability sample of schools within the catchment areas for each site were selected, and eligible youth were recruited from each school. The ABCD study approximates a multi-stage probability sample, but is not nationally-representative (46). The imaging procedures were harmonized across sites (47). The study protocol received ethics approval from [name redacted].

### Measures

#### Structural stigma

Consistent with conceptualizations of structural stigma (8) and prior research on this topic (3), we selected items that captured societal-level conditions, social/cultural norms, and/or institutional policies to create proxy measures of the social climate relevant to the three groups of interest (female, Black, and Latinx youth). We compiled items from publicly available data sources used in prior work to assess structural forms of stigma related to gender (48–50), race (51–54), and Latinx ethnicity (10). We modeled these items as indicators in a factor analysis (described below), with the final factor score determining the structural stigma score for each state for each domain of stigma (Figure 1). We chose a factor analytic approach because it 1) recognizes that different dimensions of structural stigma (e.g., norms, policies) are highly correlated; 2) improves construct validity; and 3) taps into shared variance, thereby reducing measurement error.

**Figure 1.**
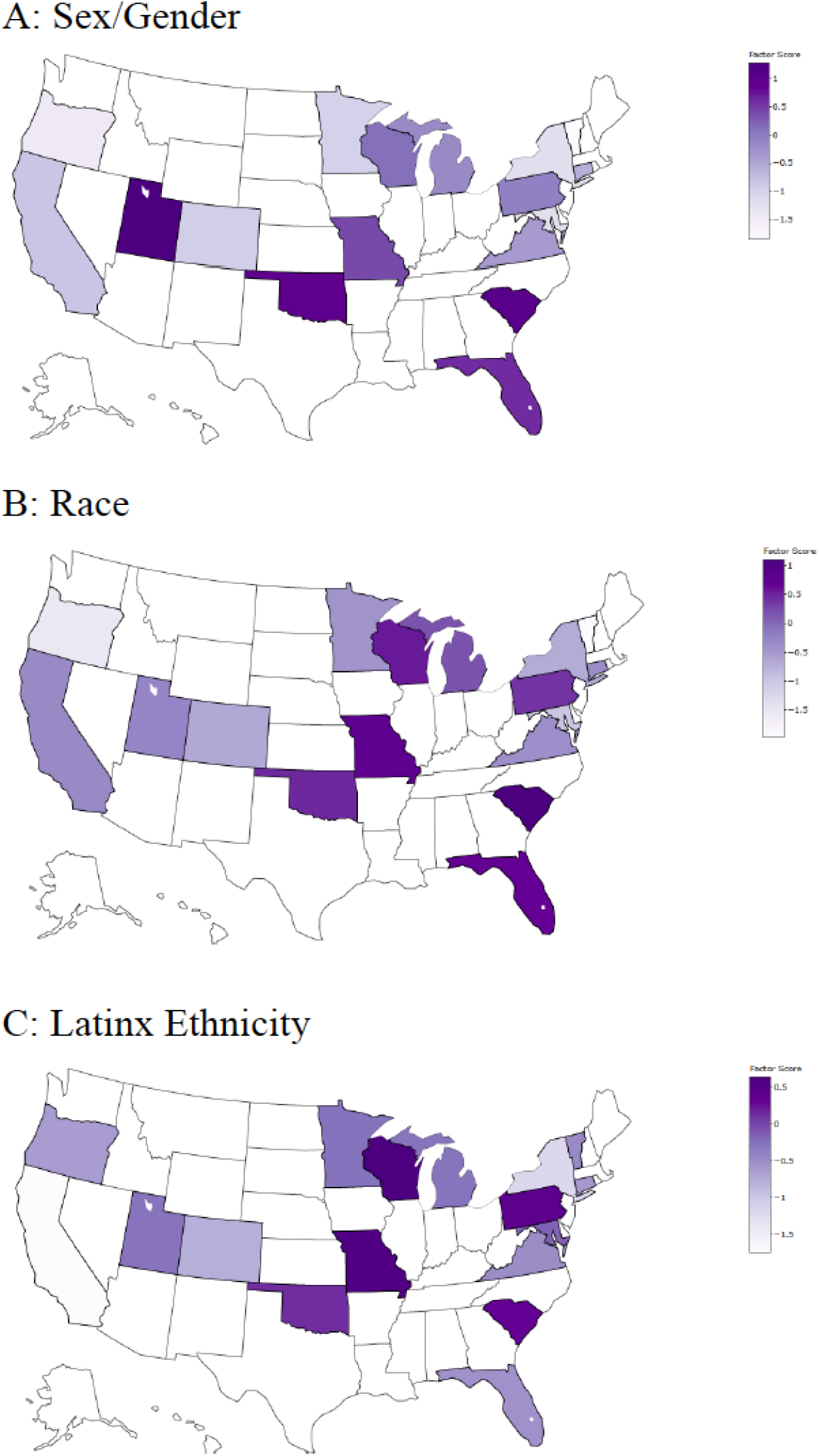
State-level structural stigma across four stigmatized groups. The figure shows the distribution of the four measures of structural stigma related to sex/gender (A), race (B), and Latinx ethnicity (C), across the 17 states in the ABCD sample.

Because we used a data-driven technique, the final factor scores included different components across the three groups. For instance, while the index of structural stigma related to Latinx ethnicity included both state laws and aggregated social norms, the index of structural stigma related to race included only aggregated norms. We nevertheless refer to all of these indicators as “structural,” in that they represent factors at the contextual rather than individual/interpersonal level. This approach is consistent with conceptualizations of individual attitudes shaping structural factors (e.g., laws and policies) in ways that subsequently influence the attitudes of individuals within a particular social context; as such, aggregated attitudes represent more than individual bias because they not only reflect but also shape broader social structures (55,56).

Below, we describe the items and sources of data for each group; further details are in the Supplement.

#### Structural Stigma Related to Gender

The measure of structural stigma related to gender was comprised of 18 items. Twelve of these indicators assessed aggregated implicit and explicit attitudes, which were obtained from two sources: Project Implicit (pooled across years 2003-2018) and the General Social Survey (pooled across years 1974-2014). The explicit indicators directly queried gender role attitudes and sexist beliefs. The implicit indicators were obtained through the Implicit Association Test (IAT) and examined to what extent respondents associate gender with career and scientific domains. The other 6 items were taken from previous state-level composite indicators of women’s social status (48,49), including: economic (e.g., ratio of men’s to women’s earnings); political (e.g., women’s representation in elected office); social and economic autonomy (e.g., women’s business ownership); and reproductive (e.g., percentage of women who live in a county without an abortion provider) factors. These items were obtained from several sources, including the Bureau of Labor Statistics, Current Population Survey, and Center for American Women in Politics.

#### Structural Stigma Related to Race

The measure of structural stigma related to race was comprised of 31 items, all of which assessed aggregated attitudes related to race and racial prejudice, which were obtained from three sources: Project Implicit (pooled across years 2002-2017), the General Social Survey (pooled across years 1973-2014), and the American National Election Survey (pooled across years 1992-2016). Collectively, these items assessed several different components of racial prejudice, including general attitudes towards Black individuals, the impact of discrimination on the lives of Black people, the existence of racial prejudice, and endorsement of racial stereotypes. A similar aggregate measure of explicit racial bias has demonstrated strong retest reliability, and convergent/discriminant validity (55).

#### Structural Stigma Related to Latinx Ethnicity

For structural stigma related to Latinx ethnicity, we used 3 indicators: 1) a feelings thermometer of explicit attitudes of immigrants (obtained from the American National Election Survey); 2) a composite index of state-level policies related to immigration (e.g., whether immigrants were granted access to health services); and 3) a feelings thermometer of explicit attitudes of Hispanics (obtained from the American National Election Survey). Our rationale for the inclusion of attitudes and policies related to immigration, despite the fact that not all Latinx individuals are immigrants, was based on previous findings (10) that immigration policies and attitudes are associated with the mental health of Latinx respondents regardless of nativity status because of: 1) the conflation of immigration with Latinx ethnicity in the US; 2) the mixed status nature of many Latinx households; 3) the concealability of immigration status, which makes people targets regardless of citizenship; and 4) the salience of immigration policy to Latinx individuals in the U.S. (57).

#### Factor analysis

We created a factor score for each state for each measure by using exploratory factor analysis with a factor loading cut off of 0.40; we reran the factor analysis iteratively and excluded variables until all retained items met the 0.40 threshold (Table S1). For each measure, a 1-factor solution emerged, indicating that these items load onto a single construct of structural stigma, providing some evidence of construct validity. Cronbach’s alpha was calculated as a measure of reliability (58). Structural stigma related to gender (α=0.94) and race (α=0.97) indicated high reliability; structural stigma related to Latinx ethnicity indicated fair reliability (α=0.57), which likely reflects the small number of items contributing to the factor.

#### Perceived discrimination

Respondents were asked a series of questions about their perceptions of discrimination, unfair treatment, and perceived acceptance based on their identity. See Supplement for items and scoring for each group. After preregistering our analysis plan, we discovered that the perceived discrimination measure was first administered in the year after the baseline neuroimaging assessment (Wave 1). The measure was then re-administered during a follow-up assessment (Wave 2), in which neuroimaging data were collected a second time. Currently, ABCD has only released half of the data on the Wave 2 sample. Because there are strengths and limitations associated with each assessment (e.g., reduced power at Wave 2 but temporal precedence, increased power at Wave 1 but lacking in temporal precedence), we present results for Wave 1 (continuous measure of perceived discrimination) in the main text and Wave 2 in the Supplement; both produce similar conclusions.

#### Brain Structure and Function

Hippocampal volume was obtained from the structural data release. Quality control measures were applied to structural MRI data, including visual inspection of structural volumes, inspection of outliers of segmented volumes for potential segmentation problems, and exclusion of data that did not meet the quality control standards in the public data release. Volume measures of left and right hippocampus, obtained using automatic segmentation in FreeSurfer 5.3, were summed to produce a measure of total hippocampal volume.

Amygdala activation to threat was measured by contrasting amygdala response to fearful faces relative to neutral faces during an emotional n-back working memory task (47). The task includes two runs of eight blocks each. On every trial, participants respond as to whether the picture was a “Match” or “No Match.” Within each block, stimuli were either all fearful faces, all neutral faces, all happy faces, or all places. Individual-level estimates of task-related BOLD signal were computed using a general linear model implemented in AFNI’s 3dDeconvolve. Hemodynamic response functions were modelled for cues (~3◻s) and trial blocks (~24◻s) as square waves convolved with a gamma variate basis function plus its temporal derivative using AFNI’s ‘SPMG’ option within 3dDeconvolve. The contrast of interest was activation during the fearful face blocks vs. activation during the neutral face blocks within an amygdala region-of-interest defined using automatic segmentation by Freesurfer 5.3. The region-of-interest coefficients for this contrast from this task were obtained from the ABCD study’s curated data release. Analyses were conducted separately for right and left amygdala, because the associations of childhood trauma with amygdala reactivity to threat often exhibit hemispheric specificity (27,29,30).

For both outcomes, ABCD guidelines were followed with regard to exclusion of participants based on data quality, motion, or inattention.

### Analytic Strategy

We pre-registered our hypotheses and analyses (see https://osf.io/9axqr). We focused on three groups of stigmatized youth in the ABCD sample: girls, Black, and Latinx youth. (In our pre-registration, we hypothesized null effects among sexual minorities at baseline, because sexual identities emerge later in development (59); see Supplement for those results.) Analyses were conducted using generalized mixed-effects models with the gamm4 function in R. Random effects included site and family. Fixed-effects included age, sex (in analyses not focused on gender), family SES (family income), parental marital status, race and ethnicity (in analyses not focused on race and Latinx ethnicity, respectively). Analyses examining hippocampal volume additionally controlled for total intracranial volume.

We discovered substantial missing data on family income (~9%). Multiple imputation (100 imputations) was used to handle this missing data. Given difficulties in imputing in 3-level multi-level models and models with small (e.g., 2 siblings) cluster sizes (60), we imputed based on 2-level structure with a random intercept of site, but not family. We also conducted supplementary analyses that controlled for an alternate measure of SES—family education— which had substantially lower missingness (0.7%) and was strongly correlated with family income (r=.64). The direction, magnitude, and significance of associations was unchanged with this alternative measure (Supplement).

A pre-registered power analyses indicated that within female participants and in all control analyses, we have adequate sample size to detect an effect size of *r*=0.1 with close to 100% power. Within Black and Latinx participants, we have sample size to detect an effect size of *r*=0.1 with 99% power.

## Results

To test our predictions, we linked the three indicators of structural to the ABCD dataset via the FIPS code of the state where each ABCD study site is located (n=21 sites) to determine their association with neural outcomes. Because our independent variable (structural stigma) is a contextual factor coded at the site-level, the degrees of freedom for the *p*-values presented below are based on the site (df=19), rather than on the total sample size of youth (e.g., n=5,489 girls) included in each analysis, which should be considered when interpreting the statistical and practical significance of the beta estimates.

### Structural Stigma and Hippocampal Volume

Figure 2 shows the results for hippocampal volume. Higher structural stigma related to gender was associated with smaller hippocampal volume among girls (n=5,489, *B*=-29.50, *SE*=16.96), although this was not statistically significant controlling for covariates (*p*=0.082).

**Figure 2.**
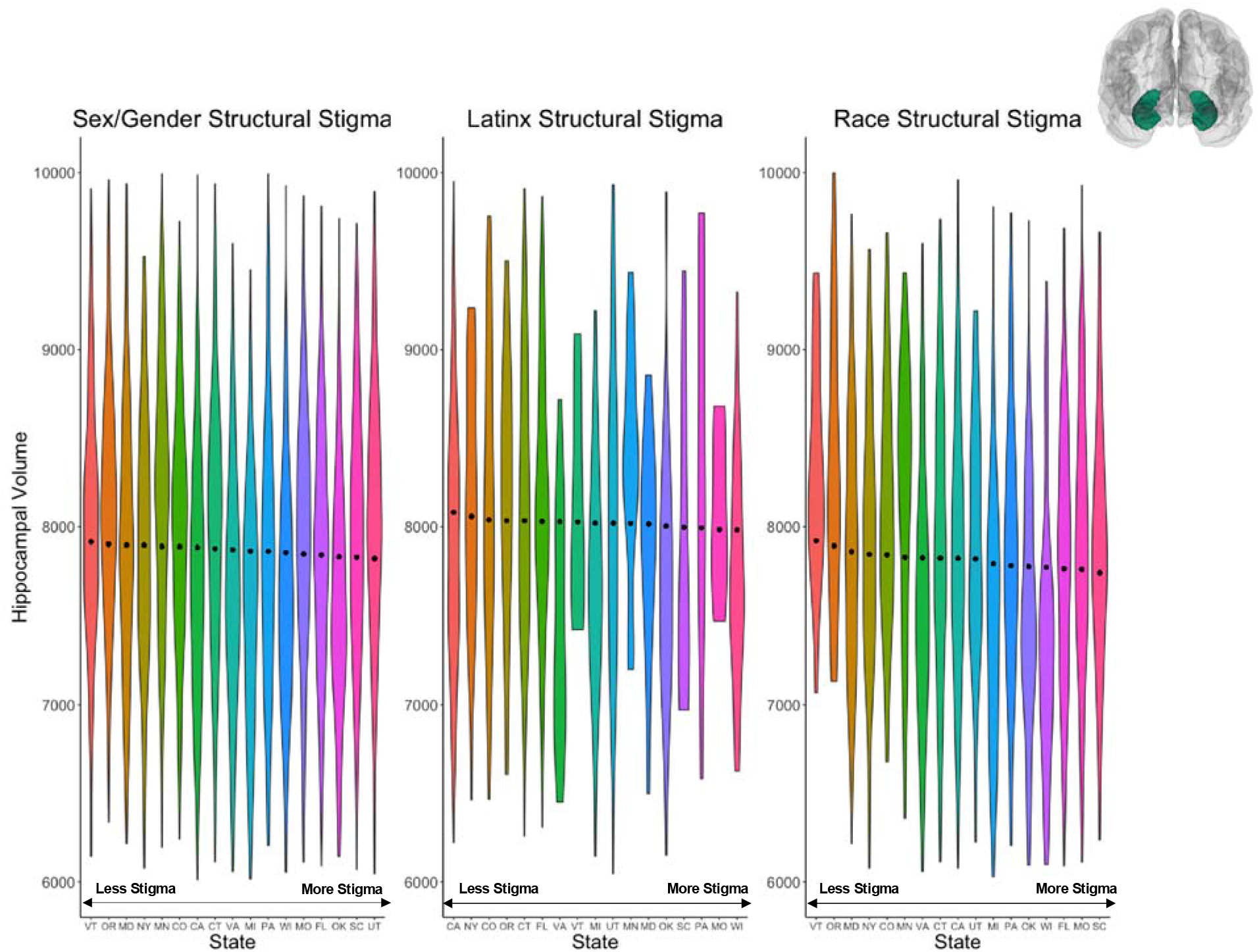
Plot of hippocampal volume among female, non-immigrant Latinx, and Black youth by state. States are ordered from left to right in each plot based on their structural stigma factor scores. Violin plots depict the density distribution of hippocampal volume in that state. Black points represent the hippocampal volume predicted based on each states’ structural stigma when all other covariates in the multilevel models of the relation between structural stigma and hippocampal volume are at their mean.

Higher structural stigma related to race was associated with smaller hippocampal volume among Black youth (n=2,421, *B*=−58.26, *SE*=25.70, *p*=0.023). A 1-unit increase in racial structural stigma was associated with a 58 mm^3^ decrease in hippocampal volume among Black youth.

Higher structural stigma related to Latinx ethnicity was associated with smaller hippocampal volume among Latinx youth (n=2,346, *B*=−40.10, *SE*=19.90, *p*=0.044). A 1-unit increase in Latinx structural stigma was associated with a 40mm^3^ decrease in hippocampal volume among Latinx youth.

As hypothesized, structural stigma was unrelated to hippocampal volume in the non-stigmatized comparison groups: boys (n=6,037, *B*=−12.7, *p*=0.486), non-Latinx White (n=6,887, *B*=6.27, *p*=0.780), and White (n=8,594, *B*=−21.56, *p*=0.231) youth.

### Structural Stigma and Amygdala Reactivity to Threat

There was no association between structural stigma and amygdala activation to fearful relative to neutral faces for any of the 3 stigmatized groups, or for the non-stigmatized comparison groups (see Supplement).

### Perceived Discrimination and Neural Outcomes

Perceived discrimination was not associated with hippocampal volume among female (*B*=−13.1, *SE*=21.5, *p*=0.542), Black (*B*=16.1, *SE*=22.5, *p*=0.473), or Latinx (*B*=−41.9, *SE*=25.5, *p*=0.101) youth. Further, no association between perceived discrimination and amygdala reactivity was observed (see Supplement).

## Discussion

We provide novel evidence that a macro-social contextual factor—structural stigma—is associated with brain structure in children, such that youth living in higher (vs. lower) structural stigma contexts had smaller hippocampal volume when they had an identity that was a target of that structural stigma. This association was observed consistently across three stigmatized groups (girls, Black, and Latinx youth), although the association reached statistical significance only for Black and Latinx youth, suggesting that this relationship is generalizable across diverse types of stigmatized identities and statuses. In contrast, we did not observe an association between structural stigma and amygdala reactivity to fearful faces. Further, perceived discrimination was not associated with neural outcomes among stigmatized youth. Thus, we provide novel evidence that an objective measure of stigma at the contextual level (structural stigma) may have a stronger influence on neurodevelopment than subjective perceptions of stigma measured at the individual level. This is the first study, to our knowledge, to simultaneously measure stigma at both structural and individual levels and to determine their relative association with neural outcomes in youth.

The associations of structural stigma with hippocampal volume build on a substantial literature documenting reduced hippocampal volume in children who have experienced trauma, who are raised in families with lower SES, and who have low levels of parental support and nurturance (25,61–63). We extend this work by demonstrating that being raised in a context characterized by higher levels of stigma towards members of one’s group similarly influences hippocampal volume as these individual-level experiences. The effect sizes for the association between structural stigma and hippocampal volume, while relatively modest in magnitude, are similar to those observed for relatively extreme stressors—like childhood trauma—that are established correlates of reduced hippocampal volume. For instance, Black youth in the highest-structural stigma states had a hippocampus that was 177 mm^3^ smaller than those in the lowest-structural stigma states, equivalent to about 2.3% of the average hippocampal volume among Black youth in the ABCD sample. A recent study, by comparison, found that the reduction in hippocampal volume attributable to childhood experiences of interpersonal violence was 364 mm^3^ (64), about 3.8% of the average volume in that sample. To further contextualize this finding, the magnitude of the observed difference in hippocampal size between high and low structural stigma states was equivalent to the predicted impact of a $20,000 difference in annual family income in this sample. Statistically small effects can have societally important consequences if they apply to many people, or if they apply repeatedly to the same person (45). These findings therefore suggest that structural stigma may be meaningfully associated with brain development in children and adolescents.

Our measure of structural stigma is a proxy for the social environment surrounding girls as well as Black and Latinx youth at the state level, which is hypothesized to influence a variety of intermediary variables that in turn may shape brain structure and function. Future research is therefore needed to identify the specific environmental and neurobiological mechanisms linking structural stigma to reduced hippocampal volume. Animal and human studies have documented lasting reductions in hippocampal volume following exposure to chronic stress (19,23–26,65,66), and as a result of low levels of support and nurturance in early life (62,63). These effects are mediated by excessive production of corticotropin-releasing hormone in animal models (19), although the precise neurobiological mechanisms contributing to these volume reductions in humans are unknown. The association of structural stigma with hippocampal volume may be due, in part, to exposure to chronic stress or lack of social support associated with living in a stigmatizing context. Stressors resulting from structural stigma are conceptualized as chronic because they are related to fairly stable underlying social structures (3). Support for a developmental pathway from structural stigma to hippocampal volume via experiences of chronic stress or low levels of support should be considered provisional, however, until it can be tested directly with longitudinal data that incorporates measures of stigma-related chronic stressors (67), which will be possible in future waves of the ABCD study. Future research is also needed to determine whether the divergent association of structural stigma with hippocampal volume and amygdala reactivity replicates and, if so, the reasons for this divergence. One possibility, supported by meta-analysis, is that task demands reduce amygdala response to salient cues (68). Because amygdala activation was assessed during a working memory task, it may have constrained variability in amygdala reactivity in our sample. The incorporation of additional tasks may help to reveal whether this contributed to the divergent neural patterns observed herein. Although our focus was on structural forms of stigma related to single identities, future research may also wish to take an intersectional approach to examining whether structural forms of stigma related to multiple axes of social stratification interact to predict neural outcomes among youth.

We note several limitations of the current study. First, this is a cross-sectional analysis. However, issues of temporality are less of a concern for causal inference in our study, given that hippocampal volume cannot cause state-level structural stigma. Further, in studies of contextual factors, one concern is whether the results are due to social selection, whereby individuals with the observed outcome (in this case, lower hippocampal volume) sort into the exposure status (i.e., structural stigma). However, in studies of children, issues of social selection are less plausible, given that children are rarely responsible for moves into and out of certain environments. Although it is possible that the selection of parents into high structural stigma environments may contribute to the observed patterns (given strong associations between family SES and hippocampal volume in children), we observed no meaningful association between structural stigma and parental SES for any group (*r*’s < 0.16). Thus, there is minimal evidence for differential selection of low-income parents into high structural stigma states.

Second, although the ABCD study is one of the largest of its kind, the study sites are located in only 17 states. This resulted in a restricted range of structural stigma for each of our groups, with more than half of the sites located in states that fell below the mean of structural stigma. This restricted range reduced our statistical power, which means that our estimates are likely conservative. At the same time, the restricted range means that different associations may be observed with another set of social contexts.

Third, our study measured structural stigma at the state level, as geographic units below the state level were not available. Our focus on distal environments offers a conservative test, given that more proximal environments are likely to exert stronger effects (69). However, this approach does not incorporate within-state heterogeneity, particularly with respect to local social environments that may differ from those at the state level. Exploring associations between structural stigma and hippocampal volume at more proximal levels of analysis (e.g., counties) therefore represents an important area for future inquiry.

Fourth, while the concept of structural stigma has existed for decades (1), its operationalization is a relatively recent occurrence (8). Existing studies have used different approaches to operationalize this construct, ranging from single-item measures of individual components of structural stigma (e.g., social attitudes (54,70–72)) to multi-item scales that capture several dimensions (48,51). Drawing on current conceptualizations of structural stigma (8), we measured structural stigma using an empirically-derived approach (i.e., factor analysis) that combined multiple indicators of societal-level conditions, social attitudes, and public policies, in order to create a comprehensive index. This approach has several advantages, including providing evidence of construct validity (i.e., showing that multiple items related to structural stigma load onto a single factor) and reducing measurement error (i.e., tapping into shared variance). However, our approach likely missed other important dimensions of structural stigma. For instance, racial disparities in incarceration, which have been used in previous studies as indicators of structural racism (51), did not load highly onto our factor. Further research is needed to determine whether these results are generalizable across different operationalizations of this construct.

Fifth, although the indicators of structural stigma were obtained across a range of years, we aggregated all responses to the state level regardless of year queried, which allowed for all states to have a sizable number of respondents, regardless of yearly sampling variation, thereby reducing measurement error. One potential limitation is that this approach does not capture changes in temporal trends in structural stigma. However, while structural sexism and structural racism have declined nationally over time, the relative levels of structural stigma in individual states (i.e., rankings relative to other states) have remained highly stable (50,73), suggesting that a time-invariant measure represents a valid approach to operationalizing this construct. Further, supplementary analyses showed that our structural stigma measures were highly correlated (*r*’s: .84 to .99) with an alternative measure that was restricted to the years following the birth of children in the ABCD sample.

An additional limitation concerns the measure of perceived discrimination. Both Black and Latinx youth were significantly more likely to endorse perceived discrimination than White and White non-Latinx youth, respectively, providing some support for construct validity. However, perceived discrimination was not higher among female than male youth. Thus, future research is needed to confirm these results with measures of perceived discrimination that have demonstrated reliability and validity.

Finally, it is possible that other contextual factors influence both the level of structural stigma as well as smaller hippocampal volume among youth. However, our negative control analyses showing no association between structural stigma and hippocampal volume for any of the non-stigmatized comparison groups help to minimize this possibility, because if structural stigma were merely a proxy for other contextual factors (e.g., state-level SES, conservatism), those other factors should operate similarly among both the stigmatized and non-stigmatized.

These limitations notwithstanding, this study not only expands our understanding of the multi-level consequences of stigma, but also suggests a potential neural mechanism underlying the established association between structural stigma and psychopathology (3). Examining whether hippocampal volume mediates the structural stigma-mental health association will be possible in future waves of the ABCD dataset as the youth age into the developmental period of risk for depression and post-traumatic stress disorder, which have both been consistently linked with hippocampal volume (32–35). In addition, our results suggest that macro-level features of the social context are associated with brain structure in children, which has implications for broadening the range of potential explanatory variables in cognitive neuroscience to include contextual influences. Collectively, these findings set the stage for future studies to identify additional contextual correlates of neurodevelopment among youth, and to uncover the environmental and neurobiological mechanisms underlying these relationships.

