## Supplemental materials for "Smaller Hippocampal Volume among Black and Latinx Youth Living in High-Stigma Contexts"

**Supplementary Information**

Creation of Independent Variable: Structural Stigma	2-10

Summary of structural stigma and participant makeup by State	11

Sensitivity Analyses with Alternate Measure of SES	12

Reproducible Pre-registered Analyses and Results	13-41

Creation of independent variable: structural stigma

The independent variable used in these analyses is state-level structural stigma, relevant to the subgroup of interest in each stratified analysis: girls, Black, and Latinx participants. Each of these variables was compiled from previously-validated sources and then modeled as indicators in a factor analysis, with the final factor score determining the structural stigma score for each state for each domain of stigma. With the exception of anti-immigrant sentiment (which had a limited inventory of possible indicators), we restricted all indicators to those that had measures for all 50 states and Washington, D.C.

The rationale for including all 50 states was to contextualize the factor scores relative to the distribution of the entire nation. Model-based factor scores are assigned based around a normal distribution centered around a mean (fixed at 0). Had we run the factor score only on states represented in the ACBD sample, estimates of structural stigma would be biased towards only our sample and may not reflect the true national mean. Our study questions were interested in measures of state-level structural stigma relative to the average of the national climate, not just the states that happened to be selected for inclusion in ABCD. After models were run using all 50 states, only the states containing ABCD sites were selected from the larger dataset. See Supplemental Table 1 for description of final variables.

All measures were coded such that higher levels corresponded to higher values of structural stigma. Measures were standardized to their average response value for all respondents, regardless of state of residence, and aggregated to the state level so that every state’s measure was the mean of the standardized individual response for respondents residing in that state. The exception was state-level variables, which were already measured at the state level and thus were not aggregated or standardized further. Measures met inclusion criteria for model selection if residents of every state and Washington, D.C., were represented in the data. The exception is anti-immigrant sentiment, which heavily relied on measures of state policies; thus, data from Washington, D.C., was not non-existent and so was not included in the models.

While the indicators often represented different survey years (e.g., General Social Survey questions ranged from the 1970’s to present day), we aggregated all responses to the state level regardless of year queried. Averaging the responses in this way allowed for all states to have a sizable number of respondents, regardless of sampling variation from year to year, thereby reducing measurement error. Further, previous analyses have shown that while structural sexism and structural racism have declined nationally over time, the relative stigma levels of individual states (i.e., rankings relative to other states) have remained stable^71,78^, suggesting that a time-invariant measure represents a valid approach to operationalizing this construct.

| **Table S1**. **Selection and Sources for Structural Stigma Variables Included in Final Factor Models** | | | |
| --- | --- | --- | --- |
| **Domain** | **Item** | **Source** | **Years available** |
| Structural Stigma Related to Sex/Gender | Tell me if you agree or disagree with this statement: Most men are better suited emotionally for politics than are most women. | General Social Survey | 1974, 1975, 1977, 1978, 1982, 1983, 1985, 1986, 1988, 1989, 1990, 1991, 1993, 1994, 1996, 1998, 2000, 2002, 2004, 2006, 2008, 2010, 2012, 2014 |
|  | It is much better for everyone involved if the man is the achiever outside the home and the woman takes care of the home and family. | General Social Survey | 1977, 1985, 1986, 1988, 1989, 1990, 1991, 1993, 1994, 1996, 1998, 2000, 2002, 2004, 2006, 2008, 2010, 2012, 2014 |
|  | Percent of women who live in a county without an abortion provider | Guttmacher Institute | 2014 |
|  | Ratio of men’s to women’s median usual weekly earnings of full-time wage and salary workers | Bureau of Labor Statistics | 2016 |
|  | Ratio of men’s to women’s labor force participation rates, age 16+ | Integrated Public Use Microdata Series Current Population Survey | 2016 |
|  | The employment and earnings composite index reflects women’s status related to occupation and income and includes four components: women’s median annual earnings, ratio of women’s to men’s earnings, women’s labor force participation, and women’s representation in managerial and professional occupations. (r) | Integrated Public Use Microdata Series; Bureau of Labor Statistics | 2013 |
|  | The political composite index reflects four components: women’s voter registration, voter turnout, representation in elected office, and the existence of institutional resources for women. The proportion of women holding political office at four levels was calculated and weighted based on prestige: (1) state representatives (weight = 1.0), (2) state senators (weight = 1.25), (3) executive officials and US representatives (weight = 1.5) (r) | Integrated Public Use Microdata Series; Center for American Women in Politics | 2015 |
|  | Social and economic autonomy composite index includes four additive components: proportion of women with health insurance, women’s educational attainment (proportion with a college degree), women’s business ownership, and proportion of women living above the federal poverty level. These were all standardized and the scores were then added together to create a composite. (r) | Integrated Public Use Microdata Series | 2013 |
|  | Prompt: *Women hold a smaller portion of the science and engineering faculty positions at top research universities than do men. The following factors are sometimes offered as reasons for this difference. Please rate how important you think each factor is for explaining this difference*  On average, whether consciously or unconsciously, men are favored in hiring and promotion. (r) | Project Implicit: gender-science | 2003-2018 |
|  | Prompt: *Women hold a smaller portion of the science and engineering faculty positions at top research universities than do men. The following factors are sometimes offered as reasons for this difference. Please rate how important you think each factor is for explaining this difference*  Directly or indirectly, boys and girls tend to receive different levels of encouragement for developing scientific interest. (r) | Project Implicit: gender-science | 2003-2018 |
|  | How strongly do you associate career and family with males and females? Career | Project Implicit: gender-career | 2005-2018 |
|  | Gender-Career Implicit Association Test: Implicit Sexism) | Project Implicit: gender-career | 2005-2018 |
|  | Gender-Science Implicit Association Test: Implicit Sexism | Project Implicit: gender-science | 2003-2018 |
|  | Prompt: *Women hold a smaller portion of the science and engineering faculty positions at top research universities than do men. The following factors are sometimes offered as reasons for this difference. Please rate how important you think each factor is for explaining this difference*  Different proportions of men and women are found among people with the very highest levels of math ability | Project Implicit: gender-science | 2003-2018 |
|  | Prompt: *Women hold a smaller portion of the science and engineering faculty positions at top research universities than do men. The following factors are sometimes offered as reasons for this difference. Please rate how important you think each factor is for explaining this difference*  On average, men and women differ in their willingness to spend time away from their families. | Project Implicit: gender-science | 2003-2018 |
|  | Prompt: *Women hold a smaller portion of the science and engineering faculty positions at top research universities than do men. The following factors are sometimes offered as reasons for this difference. Please rate how important you think each factor is for explaining this difference*  On average, men and women differ in their willingness to devote the time required by such 'high-powered' positions | Project Implicit: gender-science | 2003-2018 |
|  | Prompt: *Women hold a smaller portion of the science and engineering faculty positions at top research universities than do men. The following factors are sometimes offered as reasons for this difference. Please rate how important you think each factor is for explaining this difference*  On average, men and women differ naturally in their scientific interest | Project Implicit: gender-science | 2003-2018 |
|  | Suppose that ten men at a typical U.S. university were picked at random. How many would you predict will graduate with a scientific major (science, technology, engineering, or mathematics)? | Project Implicit: gender-science | 2003-2018 |
| Structural Stigma Related to Race | *Prompt: On the average (Negroes/Blacks/African-Americans) have worse jobs, income, and housing than white people. Do you think these differences are . . .*  Because most (Negroes/Blacks/African-Americans) just don't have the motivation or will power to pull themselves up out of poverty? | General Social Survey | 1977, 1985, 1986, 1988, 1989, 1990, 1991, 1993, 1994, 1996, 1998, 2000, 2002, 2004, 2006, 2008, 2010, 2012, 2014 |
|  | *Prompt: On the average (Negroes/Blacks/African-Americans) have worse jobs, income, and housing than white people. Do you think these differences are . . .*  Mainly due to discrimination? (r) | General Social Survey | 1977, 1985, 1986, 1988, 1989, 1990, 1991, 1993, 1994, 1996, 1998, 2000, 2002, 2004, 2006, 2008, 2010, 2012, 2014 |
|  | *Prompt: Some people think that (Blacks/Negroes/African-Americans) have been discriminated against for so long that the government has a special obligation to help improve their living standards. Others believe that the*  *government should not be giving special treatment to (Blacks/Negroes/African-Americans).*  Where would you place yourself on this scale, or haven't you made up your mind on this? | General Social Survey | 1975, 1983, 1984, 1986, 1987, 1988, 1989, 1990, 1991, 1993, 1994, 1996, 1998, 2000, 2002, 2004, 2006, 2008, 2010, 2012, 2014 |
|  | Are we spending too much, too little, or about the right amount on improving the conditions of Blacks? *[answers range from too much to too little]* (r) | General Social Survey | 1973, 1974, 1975, 1976, 1977, 1978, 1980, 1982, 1983, 1984, 1985, 1986, 1987, 1988, 1989, 1990, 1991, 1993, 1994, 1996, 1998, 2000, 2002, 2004, 2006, 2008, 2010, 2012, 2014 |
|  | *Prompt: Now I'm going to ask you about different types of contact with various groups of people. In each situation would you please tell me whether you would be very much in favor of it happening, somewhat in favor, neither in favor nor opposed to it happening, somewhat opposed, or very much opposed to it happening?*  Living in a neighborhood where half of your neighbors were blacks? | General Social Survey | 1990, 1996, 1998, 2000, 2002, 2004, 2006, 2008, 2010, 2012, 2014 |
|  | *Prompt: Now I'm going to ask you about different types of contact with various groups of people. In each situation would you please tell me whether you would be very much in favor of it happening, somewhat in favor, neither in favor nor opposed to it happening, somewhat opposed, or very much opposed to it happening?*  Having a close friend or family member marry a Black person | General Social Survey | 1990, 1996, 1998, 2000, 2002, 2004, 2006, 2008, 2010, 2012, 2014 |
|  | *Prompt: On the average (Negroes/Blacks/African-Americans) have worse jobs, income, and housing than white people. Do you think these differences are . . .*  Because most (Negroes/Blacks/African-Americans) have less in-born ability to learn? | General Social Survey | 1977, 1985, 1986, 1988, 1989, 1990, 1991, 1993, 1994, 1996, 1998, 2000, 2002, 2004, 2006, 2008, 2010, 2012, 2014 |
|  | Would you vote for a law that says a homeowner can refuse to sell to blacks, or one that says homeowners cannot refuse to sell based on skin color? | General Social Survey | 1977, 1985, 1986, 1988, 1989, 1990, 1991, 1993, 1994, 1996, 1998, 2000, 2002, 2004, 2006, 2008, 2010, 2012, 2014 |
|  | *What do you think the chances are these days that a white person won't get a job or promotion while an equally or less qualified black person gets one instead?*  Is this very likely, somewhat likely, or not very likely to happen these days? | General Social Survey | 1990, 1994, 1996, 1998, 2000, 2002, 2004, 2006, 2008, 2010, 2012, 2014 |
|  | Irish, Italians, Jewish and many other minorities overcame prejudice and worked their way up. Blacks should to the same without any special favors. 1) agree strongly; 2) agree somewhat; 3) neither agree nor disagree; 4) disagree somewhat; 5) disagree strongly (r) | ANES | 1992, 1994, 1998, 2000, 2004, 2008, 2012, 2016 |
|  | It's really a matter of some people not trying hard enough; if blacks would only try harder they could be just as well off as whites.' 1) agree strongly; 2) agree somewhat; 3) neither agree nor disagree; 4) disagree somewhat; 5) disagree strongly (r) | ANES | 1992; 1994; 2000; 2004; 2008; 2012; 2016 |
|  | If a Black person were put in charge of me, I would not mind taking advice and direction from him or her. (r) | Project-Implicit: Race | 2002-2017 |
|  | If I had a chance to introduce Black visitors to my friends and neighbors, I would be pleased to do so. (r) | Project-Implicit: Race | 2002-2017 |
|  | I would rather not have Black people live in the same apartment building I live in. | Project-Implicit: Race | 2002-2017 |
|  | I would probably feel somewhat self-conscious dancing with a Black person in a public place. | Project-Implicit: Race | 2002-2017 |
|  | I would not mind at all if a Black family with about the same income and education as me moved in next door. (r) | Project-Implicit: Race | 2002-2017 |
|  | I think that Black people look more similar to each other than White people do. | Project-Implicit: Race | 2002-2017 |
|  | Interracial marriage should be discouraged to avoid the 'who-am-I?' confusion that the children feel. | Project-Implicit: Race | 2002-2017 |
|  | I get very upset when I hear a White person make a prejudicial remark about Black people. (r) | Project-Implicit: Race | 2002-2017 |
|  | I favor open housing laws that allow more racial integration of neighborhoods. (r) | Project-Implicit: Race | 2002-2017 |
|  | It would not bother me if my new roommate was Black. (r) | Project-Implicit: Race | 2002-2017 |
|  | It is likely that Black people will bring violence to neighborhoods when they move in. | Project-Implicit: Race | 2002-2017 |
|  | The federal government should take decisive steps to override the injustices Black people suffer at the hands of local authorities. (r) | Project-Implicit: Race | 2002-2017 |
|  | Black and White people are inherently equal. (r) | Project-Implicit: Race | 2002-2017 |
|  | Black people are demanding too much too fast in their push for equal rights. | Project-Implicit: Race | 2002-2017 |
|  | White people should support Black people in their struggle against discrimination and segregation. (r) | Project-Implicit: Race | 2002-2017 |
|  | Generally, Black people are not as smart as White people. | Project-Implicit: Race | 2002-2017 |
|  | I worry that in the next few years I may be denied my application for a job or a promotion because of preferential treatment given to minority group members. | Project-Implicit: Race | 2002-2017 |
|  | Some Black people are so touchy about race that it is difficult to get along with them. | Project-Implicit: Race | 2002-2017 |
|  | Do Black people tend to be violence prone or do they tend not to be prone to violence? (r) | Project-Implicit: Race | 2002-2017 |
|  | Do Black people tend to prefer to be self-supporting or do they tend to prefer to live off welfare? | Project-Implicit: Race | 2002-2017 |
| Structural Stigma Related to Latinx Ethnicity | Illegal immigrants feelings thermometer (r) | ANES | 2004, 2008, 2012, 2016 |
|  | Hispanics feelings thermometer (r) | ANES | 1996, 2000, 2002, 2004, 2008, 2012, 2016 |
|  | Index for the following items: Access to health services for immigrants; Cooperation by state or local law enforcement with federal immigration enforcement; Use of non-English language; Immigrant employment in a broad segment of the private sector; Restricts immigrant access to a broad class of business licenses; Access to rental housing for immigrants; Prohibits discrimination based on citizenship or immigration status; Providing identification to access relevant services or opportunities; Access to higher education for immigrants; Access to driver’s licenses for immigrants (r) | Wake Forest | 2016 |

Notes: (r) indicates items that were reverse scored so that higher ratings represented higher levels

of structural sexism. ANES=American National Election Survey.

|  |  | Female youth | | | Black youth | | | Latinx youth | | | LGB youth | | |
| --- | --- | --- | --- | --- | --- | --- | --- | --- | --- | --- | --- | --- | --- |
| State | Total | % of total | SS Factor | SS Rank | % of total | SS Factor | SS Rank | % of total | SS Factor | SS Rank | % of total | SS Factor | SS Rank |
| CA | 1800 | 48% | -0.83 | 11 | 10% | -0.31 | 9 | 50% | -1.75 | 17 | 0.17% | -1.40 | 17 |
| CO | 554 | 47% | -0.93 | 12 | 9% | -0.65 | 13 | 22% | -0.76 | 15 | 0.72% | -0.79 | 12 |
| CT | 620 | 46% | -0.57 | 10 | 28% | -0.33 | 10 | 23% | -0.59 | 13 | 0.16% | -0.83 | 13 |
| FL | 1075 | 48% | 0.61 | 4 | 24% | 0.70 | 3 | 46% | -0.53 | 12 | 0.19% | -0.13 | 8 |
| MD | 591 | 48% | -1.20 | 15 | 46% | -0.96 | 15 | 7% | -0.16 | 6 | 1.02% | -0.62 | 11 |
| MI | 706 | 48% | -0.15 | 8 | 27% | 0.20 | 7 | 9% | -0.28 | 9 | 0.14% | 0.19 | 6 |
| MN | 595 | 46% | -0.98 | 13 | 8% | -0.44 | 12 | 6% | -0.25 | 7 | 0.17% | -0.37 | 10 |
| MO | 684 | 49% | 0.41 | 5 | 37% | 0.76 | 2 | 4% | 0.59 | 2 | 0.29% | 0.41 | 4 |
| NY | 339 | 45% | -1.19 | 14 | 29% | -0.68 | 14 | 14% | -1.20 | 16 | 0.29% | -1.28 | 15 |
| OK | 734 | 48% | 0.96 | 3 | 24% | 0.49 | 5 | 19% | 0.13 | 5 | 0.27% | 1.10 | 1 |
| OR | 582 | 50% | -1.44 | 16 | 8% | -1.52 | 16 | 16% | -0.62 | 14 | 0.69% | -1.17 | 14 |
| PA | 427 | 46% | -0.05 | 7 | 63% | 0.40 | 6 | 4% | 0.39 | 3 | 0.23% | -0.03 | 7 |
| SC | 375 | 50% | 1.05 | 2 | 34% | 1.11 | 1 | 4% | 0.32 | 4 | 0.00% | 1.05 | 2 |
| UT | 1000 | 44% | 1.28 | 1 | 3% | -0.29 | 8 | 11% | -0.26 | 8 | 0.40% | 1.04 | 3 |
| VA | 538 | 51% | -0.37 | 9 | 35% | -0.37 | 11 | 8% | -0.49 | 11 | 0.56% | 0.32 | 5 |
| VT | 573 | 48% | -1.85 | 17 | 3% | -1.98 | 17 | 3% | -0.45 | 10 | 0.17% | -1.38 | 16 |
| WI | 340 | 44% | 0.13 | 6 | 17% | 0.58 | 4 | 10% | 0.64 | 1 | 0.59% | -0.16 | 9 |
| Note: Total = total number of participants from that state included in analyses of hippocampal volume. SS= structural stigma | | | | | | | | | | | | | |

**Table S2: Summary of structural stigma and participant makeup by State.**

Sensitivity Analyses with Alternate Measure of SES (Parental Education)

As mentioned in the main text, there was a large amount of missingness (~9%) of parental income. Thus, we also conducted supplementary analyses that instead controlled for family education, which had substantially lower missingness (0.7%) and was strongly correlated with family income (*r*=.64). These results were of the same direction and magnitude as those reported in the main text: Controlling for study covariates, higher structural stigma related to sex/gender was marginally associated with smaller hippocampal volume among girls (n=5,366, *B*=-31.18, *SE*=16.94, *p*=0.066); a 1-unit increase in sex/gender structural stigma was associated with a 31 mm^3^ decrease in hippocampal volume among girls. Similarly, higher structural stigma related to race was associated with smaller hippocampal volume among Black youth (n=2,314, *B*=-61.36, *SE*=25.49, *p*=0.016); a 1-unit increase in racial structural stigma was associated with a 61 mm^3^ decrease in hippocampal volume among Black youth. Higher structural stigma related to Latinx ethnicity was associated with smaller hippocampal volume among Latinx youth (n=2,284, *B*=-45.47, *SE*=21.55, *p*=0.035); a 1-unit increase in Latinx structural stigma was associated with a 45mm^2^ decrease in hippocampal volume among Latinx youth. In a pre-specified sensitivity analysis, we found that structural stigma related to Latinx ethnicity was not associated with hippocampal volume among Latinx youth who were born outside the U.S. or who had parents who were born outside the U.S (n=1,319, *B*=16.61, *SE*=38.06, *p*=0.663); rather, the association between structural stigma and smaller hippocampal volume was only observed among U.S-born Latinx youth (n=965, *B*=-87.64, *SE*=28.93, *p*=0.003).

Reproducible Pre-registered Analyses and Results

### Preregistered analyses

The following analyses were preregistered at on <https://osf.io/9axqr> on January 27, 2020.

### Analyses involving Structural Stigma

H1.1a IV = Sexual orientation structural stigma; DVs = amygdala activation to negative vs. neutral faces, separately for right and left amygdala (tfmri_nback_all_224, tfmri_nback_all_238); covariates = age, sex at birth, marital status of household, parental education, race, and ethnicity; expected association = null

```
## Primary Analyses
lamod <- hipvol1 ~ sexual_orientation_factor + sex + race.white + latinx + unmarried +
    income ~ 1 + (1 | site_id_l)
lgbmodla <- gamm4(Lamyg1 ~ sexual_orientation_factor + sex + race.white + unmarried +
    income, random = ~(1 | site_id_l), data = lgbc_amyg1)
lgbmodra <- gamm4(Ramyg1 ~ sexual_orientation_factor + sex + race.white + unmarried +
    income, random = ~(1 | site_id_l), data = lgbc_amyg1)
# Control Analyses in non-stigmatized group
la.imp2 <- panImpute(ss_amyg[which(ss_amyg$lgbc == "No"), ], formula = lamod, m = 100,
    seed = 123, silent = TRUE)
laList2 <- mitmlComplete(la.imp2, print = "all")
lgbmodla2 <- with(laList2, lmer(Lamyg1 ~ sexual_orientation_factor + sex + race.white +
    latinx + unmarried + income + (1 | site_id_l)))
lgbmodra2 <- with(laList2, lmer(Ramyg1 ~ sexual_orientation_factor + sex + race.white +
    latinx + unmarried + income + (1 | site_id_l)))
# Sensitivity analysis where sexual minority is defined by child report of 'yes'
# or 'maybe'
la.imp3 <- panImpute(lgbc_amyg2, formula = lamod, m = 100, seed = 123, silent = TRUE)
laList3 <- mitmlComplete(la.imp3, print = "all")
lgbmodla3 <- with(laList3, lmer(Lamyg1 ~ sexual_orientation_factor + sex + race.white +
    latinx + unmarried + income + (1 | site_id_l)))
lgbmodra3 <- with(laList3, lmer(Ramyg1 ~ sexual_orientation_factor + sex + race.white +
    latinx + unmarried + income + (1 | site_id_l)))
# Sensitivity analysis where sexual minority is defined by parent report of 'yes'
# or 'maybe'
la.imp4 <- panImpute(lgbp_amyg, formula = lamod, m = 100, seed = 123, silent = TRUE)
laList4 <- mitmlComplete(la.imp4, print = "all")
lgbmodla4 <- with(laList4, lmer(Lamyg1 ~ sexual_orientation_factor + sex + race.white +
    latinx + unmarried + income + (1 | site_id_l)))
lgbmodra4 <- with(laList4, lmer(Ramyg1 ~ sexual_orientation_factor + sex + race.white +
    latinx + unmarried + income + (1 | site_id_l)))
## Primary Analyses
summary(lgbmodla$gam)
##
## Family: gaussian
## Link function: identity
##
## Formula:
## Lamyg1 ~ sexual_orientation_factor + sex + race.white + unmarried +
##     income
##
## Parametric coefficients:
##                             Estimate Std. Error t value Pr(>|t|)
## (Intercept)               -1.755e-01  2.660e-01  -0.660    0.515
## sexual_orientation_factor  8.260e-02  1.269e-01   0.651    0.521
## sexM                       4.649e-04  1.374e-01   0.003    0.997
## race.white                 9.328e-02  2.043e-01   0.457    0.652
## unmarried                  1.676e-02  2.205e-01   0.076    0.940
## income                     7.543e-07  1.445e-06   0.522    0.606
##
##
## R-sq.(adj) =  -0.155
## lmer.REML = 63.537  Scale est. = 0.10991   n = 32
summary(lgbmodra$gam)
##
## Family: gaussian
## Link function: identity
##
## Formula:
## Ramyg1 ~ sexual_orientation_factor + sex + race.white + unmarried +
##     income
##
## Parametric coefficients:
##                             Estimate Std. Error t value Pr(>|t|)
## (Intercept)               -6.831e-01  4.496e-01  -1.519    0.141
## sexual_orientation_factor -1.138e-01  1.538e-01  -0.740    0.466
## sexM                       2.075e-01  2.451e-01   0.847    0.405
## race.white                 5.276e-01  3.421e-01   1.542    0.135
## unmarried                  3.907e-01  3.738e-01   1.045    0.306
## income                     2.194e-06  2.517e-06   0.871    0.391
##
##
## R-sq.(adj) =  0.0183
## lmer.REML = 88.899  Scale est. = 0.43468   n = 32
# Control Analyses in non-stigmatized group
testEstimates(lgbmodla2)$estimates[, 1:5]
##                                Estimate    Std.Error    t.value           df
## (Intercept)                9.873632e-02 3.086265e-02  3.1992171 1.142838e+05
## sexual_orientation_factor  5.153470e-03 1.069459e-02  0.4818763 1.091358e+07
## sexM                      -2.314933e-02 1.791734e-02 -1.2920072 1.396845e+11
## race.white                -3.131928e-02 2.300527e-02 -1.3613958 5.441076e+06
## latinx                    -1.347210e-02 2.375406e-02 -0.5671493 2.632662e+06
## unmarried                  5.129405e-03 2.306351e-02  0.2224034 2.276161e+05
## income                    -7.630032e-08 1.864423e-07 -0.4092436 1.614606e+04
##                               P(>|t|)
## (Intercept)               0.001378389
## sexual_orientation_factor 0.629893829
## sexM                      0.196354656
## race.white                0.173388689
## latinx                    0.570612830
## unmarried                 0.824000033
## income                    0.682366365
testEstimates(lgbmodra2)$estimates[, 1:5]
##                                Estimate    Std.Error    t.value           df
## (Intercept)                9.484432e-02 2.912193e-02  3.2568009 8.347051e+04
## sexual_orientation_factor  1.436816e-02 1.006988e-02  1.4268454 6.468973e+06
## sexM                      -1.506668e-02 1.686313e-02 -0.8934687 1.032431e+11
## race.white                -2.024592e-02 2.165889e-02 -0.9347626 4.072883e+06
## latinx                     3.162048e-03 2.242489e-02  0.1410062 6.657764e+05
## unmarried                 -1.074206e-02 2.173240e-02 -0.4942877 1.840869e+05
## income                    -2.516649e-08 1.766348e-07 -0.1424776 1.211514e+04
##                               P(>|t|)
## (Intercept)               0.001127204
## sexual_orientation_factor 0.153624497
## sexM                      0.371606212
## race.white                0.349910717
## latinx                    0.887865091
## unmarried                 0.621103615
## income                    0.886705180
# Sensitivity analysis where sexual minority is defined by child report of 'yes'
# or 'maybe'
testEstimates(lgbmodla3)$estimates[, 1:5]
##                                Estimate    Std.Error    t.value           df
## (Intercept)                5.681270e-02 2.110314e-01  0.2692145    12291.277
## sexual_orientation_factor -8.537655e-02 1.072942e-01 -0.7957237   337708.721
## sexM                       1.844388e-01 1.102844e-01  1.6723923 54523459.250
## race.white                 3.002662e-02 1.604364e-01  0.1871559   143762.610
## latinx                    -1.112998e-01 1.742010e-01 -0.6389159   315898.344
## unmarried                 -1.747175e-01 1.581473e-01 -1.1047767     9456.071
## income                     1.834244e-07 1.343648e-06  0.1365123     1264.578
##                              P(>|t|)
## (Intercept)               0.78776912
## sexual_orientation_factor 0.42619321
## sexM                      0.09444699
## race.white                0.85153868
## latinx                    0.52287808
## unmarried                 0.26928447
## income                    0.89143806
testEstimates(lgbmodra3)$estimates[, 1:5]
##                                Estimate    Std.Error    t.value          df
## (Intercept)               -3.523214e-02 1.848032e-01 -0.1906468   15296.568
## sexual_orientation_factor -5.232091e-02 6.609426e-02 -0.7916105   80179.260
## sexM                       6.202268e-02 1.055482e-01  0.5876244 6688196.554
## race.white                 1.337243e-01 1.480559e-01  0.9032011  526748.961
## latinx                     2.104387e-01 1.576253e-01  1.3350568   60350.098
## unmarried                 -4.114223e-02 1.458828e-01 -0.2820225   20037.191
## income                    -2.421463e-07 1.203022e-06 -0.2012817    2097.502
##                             P(>|t|)
## (Intercept)               0.8488049
## sexual_orientation_factor 0.4285902
## sexM                      0.5567845
## race.white                0.3664196
## latinx                    0.1818628
## unmarried                 0.7779292
## income                    0.8404979
# Sensitivity analysis where sexual minority is defined by parent report of 'yes'
# or 'maybe'
testEstimates(lgbmodla4)$estimates[, 1:5]
##                                Estimate    Std.Error     t.value           df
## (Intercept)                2.468950e-02 9.335772e-02  0.26446119 2.070188e+06
## sexual_orientation_factor  1.881893e-02 2.526802e-02  0.74477263 6.820190e+07
## sexM                       1.053099e-01 4.131952e-02  2.54867191 3.123336e+10
## race.white                 8.884517e-03 7.661107e-02  0.11596911 3.294044e+08
## latinx                    -8.071445e-02 6.425511e-02 -1.25615611 6.936716e+06
## unmarried                 -5.021566e-03 5.413548e-02 -0.09275923 3.072909e+06
## income                    -6.133541e-08 4.222663e-07 -0.14525292 9.833916e+04
##                             P(>|t|)
## (Intercept)               0.7914246
## sexual_orientation_factor 0.4564092
## sexM                      0.0108134
## race.white                0.9076770
## latinx                    0.2090594
## unmarried                 0.9260948
## income                    0.8845115
testEstimates(lgbmodra4)$estimates[, 1:5]
##                                Estimate    Std.Error     t.value           df
## (Intercept)                1.049507e-01 8.915112e-02  1.17722226    2228706.4
## sexual_orientation_factor  3.002928e-03 2.413200e-02  0.12443758   73401475.5
## sexM                       4.942782e-02 3.946371e-02  1.25248782 8565879400.0
## race.white                -1.708402e-02 7.316844e-02 -0.23348898  325968979.6
## latinx                    -6.654031e-02 6.139407e-02 -1.08382303    4606261.9
## unmarried                 -1.281983e-03 5.168373e-02 -0.02480437    4049235.1
## income                    -3.858454e-07 4.023943e-07 -0.95887379     131754.5
##                             P(>|t|)
## (Intercept)               0.2391069
## sexual_orientation_factor 0.9009688
## sexM                      0.2103922
## race.white                0.8153817
## latinx                    0.2784433
## unmarried                 0.9802110
## income                    0.3376241
```

**Result: Sexual orientation structural stigma was not significantly associated with amygdala reactivity among youth who self-identified as LGB.**

H1.2a IV = Sexual orientation structural stigma; DVs = bilateral hippocampal volume, calculated by taking the mean volume across the right and left hemispheres (smri_vol_scs_hpuslh, smri_vol_scs_hpusrh); covariates = total intracranial volume, age, sex at birth, marital status of household, parental education, race, and ethnicity; expected association = null

```
## Primary analyses Multiple imputation
lhmod <- sexual_orientation_factor + sex + race.white + latinx + intracranialv1 +
    unmarried + income ~ 1 + (1 | site_id_l)
lh.imp <- panImpute(lgbc_hip1, formula = lhmod, m = 100, seed = 123, silent = TRUE)
lhList <- mitmlComplete(lh.imp, print = "all")
# Primary Analysis
lgbmod <- with(lhList, lmer(hipvol1 ~ sexual_orientation_factor + sex + race.white +
    latinx + intracranialv1 + unmarried + income + (1 | site_id_l)))
# Control Analysis in non-stigmatized group
lh.imp2 <- panImpute(ss_hip[which(ss_hip$lgbc == "No"), ], formula = lhmod, m = 100,
    seed = 123, silent = TRUE)
lhList2 <- mitmlComplete(lh.imp2, print = "all")
lgbmod2 <- with(lhList2, lmer(hipvol1 ~ sexual_orientation_factor + sex + race.white +
    latinx + intracranialv1 + unmarried + income + (1 | site_id_l/rel_family_id)))
# Sensitivity analysis where sexual minority is defined by child report of 'yes'
# or 'maybe'
lh.imp3 <- panImpute(lgbc_hip2, formula = lhmod, m = 100, seed = 123, silent = TRUE)
lhList3 <- mitmlComplete(lh.imp3, print = "all")
lgbmod3 <- with(lhList3, lmer(hipvol1 ~ sexual_orientation_factor + sex + race.white +
    latinx + intracranialv1 + unmarried + income + (1 | site_id_l)))
# Sensitivity analysis where sexual minority is defined by parent report of 'yes'
# or 'maybe'
lh.imp4 <- panImpute(lgbp_hip, formula = lhmod, m = 100, seed = 123, silent = TRUE)
lhList4 <- mitmlComplete(lh.imp4, print = "all")
lgbmod4 <- with(lhList4, lmer(hipvol1 ~ sexual_orientation_factor + sex + race.white +
    latinx + intracranialv1 + unmarried + income + (1 | site_id_l)))
# Primary Analysis
testEstimates(lgbmod)$estimates[, 1:5]
##                                Estimate    Std.Error      t.value           df
## (Intercept)               -1.248403e+03 9.172102e+02 -1.361087679  89498397.83
## sexual_orientation_factor  7.359533e+01 9.980896e+01  0.737362026  11677186.74
## sexM                      -5.625127e+02 1.840582e+02 -3.056166955   7307096.92
## race.white                 6.672093e+02 1.955865e+02  3.411325694 217087923.14
## latinx                     4.284056e+02 2.354463e+02  1.819546571   3596820.45
## intracranialv1             5.777389e-03 6.350758e-04  9.097163772   1968957.95
## unmarried                  2.632998e+02 2.194931e+02  1.199581213     94149.70
## income                     3.233983e-06 1.683669e-03  0.001920795     14911.69
##                                P(>|t|)
## (Intercept)               0.1734859864
## sexual_orientation_factor 0.4609022364
## sexM                      0.0022418719
## race.white                0.0006464783
## latinx                    0.0688281676
## intracranialv1            0.0000000000
## unmarried                 0.2303050451
## income                    0.9984674536
# Control Analysis in non-stigmatized group
testEstimates(lgbmod2)$estimates[, 1:5]
##                                Estimate    Std.Error   t.value           df
## (Intercept)                2.612694e+03 8.477725e+01 30.818331 5.623303e+07
## sexual_orientation_factor -2.144813e+01 1.898347e+01 -1.129832 8.421870e+07
## sexM                       5.134572e+01 1.533699e+01  3.347835 1.731833e+08
## race.white                 8.994545e+01 1.796872e+01  5.005670 2.110266e+06
## latinx                     2.603237e+01 2.025250e+01  1.285390 2.627626e+05
## intracranialv1             3.533193e-03 5.762959e-05 61.308661 4.883770e+07
## unmarried                 -2.616901e+01 1.780018e+01 -1.470154 1.261460e+05
## income                     7.727005e-04 1.497819e-04  5.158837 8.428717e+03
##                                P(>|t|)
## (Intercept)               0.000000e+00
## sexual_orientation_factor 2.585472e-01
## sexM                      8.144557e-04
## race.white                5.567263e-07
## latinx                    1.986570e-01
## intracranialv1            0.000000e+00
## unmarried                 1.415225e-01
## income                    2.541478e-07
# Sensitivity analysis where sexual minority is defined by child report of 'yes'
# or 'maybe'
testEstimates(lgbmod3)$estimates[, 1:5]
##                                Estimate    Std.Error      t.value           df
## (Intercept)               -1.248403e+03 9.172102e+02 -1.361087679  89498397.83
## sexual_orientation_factor  7.359533e+01 9.980896e+01  0.737362026  11677186.74
## sexM                      -5.625127e+02 1.840582e+02 -3.056166955   7307096.92
## race.white                 6.672093e+02 1.955865e+02  3.411325694 217087923.14
## latinx                     4.284056e+02 2.354463e+02  1.819546571   3596820.45
## intracranialv1             5.777389e-03 6.350758e-04  9.097163772   1968957.95
## unmarried                  2.632998e+02 2.194931e+02  1.199581213     94149.70
## income                     3.233983e-06 1.683669e-03  0.001920795     14911.69
##                                P(>|t|)
## (Intercept)               0.1734859864
## sexual_orientation_factor 0.4609022364
## sexM                      0.0022418719
## race.white                0.0006464783
## latinx                    0.0688281676
## intracranialv1            0.0000000000
## unmarried                 0.2303050451
## income                    0.9984674536
# Sensitivity analysis where sexual minority is defined by parent report of 'yes'
# or 'maybe'
testEstimates(lgbmod4)$estimates[, 1:5]
##                                Estimate    Std.Error    t.value           df
## (Intercept)                3.096635e+03 2.621426e+02 11.8127883 8.985885e+08
## sexual_orientation_factor -4.254107e+01 4.210247e+01 -1.0104174 1.372010e+08
## sexM                       3.117364e+01 4.925265e+01  0.6329333 3.102673e+09
## race.white                 2.231732e+02 7.570660e+01  2.9478698 1.195113e+08
## latinx                    -2.935152e+01 6.649691e+01 -0.4413968 8.943492e+05
## intracranialv1             3.194417e-03 1.736139e-04 18.3995441 2.621485e+08
## unmarried                 -4.888380e+01 5.655976e+01 -0.8642858 1.446266e+06
## income                     3.308770e-04 4.476163e-04  0.7391979 6.179097e+04
##                               P(>|t|)
## (Intercept)               0.000000000
## sexual_orientation_factor 0.312295370
## sexM                      0.526777174
## race.white                0.003199718
## latinx                    0.658925848
## intracranialv1            0.000000000
## unmarried                 0.387431063
## income                    0.459789657
```

**Result: Sexual orientation structural stigma was not significantly associated with hippocampal volume among youth who self-identified as LGB.**

H1.3a IV = Sex/gender Structural stigma; DVs = amygdala activation to negative vs. neutral faces, separately for right and left amygdala (tfmri_nback_all_224, tfmri_nback_all_238); covariates = age, marital status of household, parental education, race, and ethnicity; expected association = positive

```
## Primary Analyses Multiple imputation
samod <- sexism_factor + race.white + latinx + unmarried + income ~ 1 + (1 | site_id_l)
sa.imp <- panImpute(ss_amyg[which(ss_amyg$sex == "F"), ], formula = samod, m = 100,
    seed = 123, silent = TRUE)
saList <- mitmlComplete(sa.imp, print = "all")
# Primary analysis Left
sexismmodla <- with(saList, lmer(Lamyg1 ~ sexism_factor + race.white + latinx + unmarried +
    income + (1 | site_id_l/rel_family_id)))
# Right
sexismmodra <- with(saList, lmer(Ramyg1 ~ sexism_factor + race.white + latinx + unmarried +
    income + (1 | site_id_l/rel_family_id)))

# Control Analyses in non-stigmatized group
sa.imp2 <- panImpute(ss_amyg[which(ss_amyg$sex == "M"), ], formula = samod, m = 100,
    seed = 123, silent = TRUE)
saList2 <- mitmlComplete(sa.imp2, print = "all")
sexismmodla2 <- with(saList2, lmer(Lamyg1 ~ sexism_factor + race.white + latinx +
    unmarried + income + (1 | site_id_l/rel_family_id)))
sexismmodra2 <- with(saList2, lmer(Ramyg1 ~ sexism_factor + race.white + latinx +
    unmarried + income + (1 | site_id_l/rel_family_id)))
## Primary Analyses
testEstimates(sexismmodla)$estimates[, 1:5]
##                    Estimate    Std.Error     t.value          df    P(>|t|)
## (Intercept)    3.640172e-02 3.578514e-02  1.01723000   226317.36 0.30904511
## sexism_factor -2.082853e-04 1.220144e-02 -0.01707055 30054010.37 0.98638034
## race.white    -3.547650e-02 2.828457e-02 -1.25427074  9958161.24 0.20974364
## latinx         5.381325e-03 2.855750e-02  0.18843827  2478479.33 0.85053312
## unmarried      1.776079e-02 2.858501e-02  0.62133231   809943.66 0.53438118
## income         3.819484e-07 2.262996e-07  1.68779969    34049.69 0.09145884
testEstimates(sexismmodra)$estimates[, 1:5]
##                    Estimate    Std.Error    t.value          df   P(>|t|)
## (Intercept)    4.497134e-02 3.214000e-02  1.3992327   319146.13 0.1617442
## sexism_factor  4.760472e-03 1.097524e-02  0.4337465 51896780.37 0.6644725
## race.white    -1.484253e-02 2.543861e-02 -0.5834645 16363848.25 0.5595806
## latinx         2.810847e-02 2.573465e-02  1.0922421  1195675.68 0.2747269
## unmarried      1.425894e-02 2.570472e-02  0.5547207   973584.01 0.5790859
## income         2.653162e-07 2.025684e-07  1.3097612    50358.31 0.1902826
# Control Analyses in non-stigmatized group
testEstimates(sexismmodla2)$estimates[, 1:5]
##                    Estimate    Std.Error    t.value          df     P(>|t|)
## (Intercept)    9.835308e-02 3.389728e-02  2.9015037   30348.811 0.003716433
## sexism_factor  6.476297e-03 1.108427e-02  0.5842783 1709425.448 0.559033181
## race.white    -1.846209e-02 2.680358e-02 -0.6887920 1445838.963 0.490954293
## latinx        -1.867650e-02 2.627077e-02 -0.7109232 1289558.460 0.477131978
## unmarried     -4.230132e-02 2.677152e-02 -1.5800865   66283.910 0.114091823
## income        -2.388219e-07 2.187551e-07 -1.0917318    4977.826 0.275003834
testEstimates(sexismmodra2)$estimates[, 1:5]
##                    Estimate    Std.Error    t.value          df    P(>|t|)
## (Intercept)    9.148499e-02 3.256036e-02  2.8097045   27744.575 0.00496217
## sexism_factor  4.983658e-03 1.063734e-02  0.4685061 1706354.217 0.63942274
## race.white    -1.190754e-02 2.568004e-02 -0.4636888 1790306.127 0.64287079
## latinx        -1.295210e-02 2.521258e-02 -0.5137158  851075.668 0.60745080
## unmarried     -4.217444e-02 2.573619e-02 -1.6387212   59416.029 0.10127663
## income        -1.225897e-07 2.101406e-07 -0.5833700    4920.139 0.55967099
```

**Result: Sex/gender structural stigma was not significantly associated with amygdala reactivity among female youth**

H1.4a IV = Sex/gender Structural stigma; DVs = bilateral hippocampal volume, calculated by taking the mean volume across the right and left hemispheres (smri_vol_scs_hpuslh, smri_vol_scs_hpusrh); covariates = total intracranial volume, age, marital status of household, parental education, race, and ethnicity; expected association = negative

```
## Primary analyses Multiple imputation
shmod <- sexism_factor + race.white + latinx + intracranialv1 + unmarried + income ~
    1 + (1 | site_id_l)
sh.imp <- panImpute(sexism_hip, formula = shmod, m = 100, seed = 123, silent = TRUE)
shList <- mitmlComplete(sh.imp, print = "all")
# Primary analysis
sexismmod <- with(shList, lmer(hipvol1 ~ sexism_factor + race.white + latinx + intracranialv1 +
    unmarried + income + (1 | site_id_l/rel_family_id)))
# Control Analyis in non-stigmatized group
sh.imp2 <- panImpute(ss_hip[which(ss_hip$sex == "M"), ], formula = shmod, m = 100,
    seed = 123, silent = TRUE)
shList2 <- mitmlComplete(sh.imp2, print = "all")
sexismmod2 <- with(shList2, lmer(hipvol1 ~ sexism_factor + race.white + latinx +
    intracranialv1 + unmarried + income + (1 | site_id_l/rel_family_id)))
# Primary analysis
testEstimates(sexismmod)$estimates[, 1:5]
##                     Estimate    Std.Error    t.value           df      P(>|t|)
## (Intercept)     2.417638e+03 1.041571e+02 23.2114631 1.150698e+08 0.000000e+00
## sexism_factor  -2.950117e+01 1.696274e+01 -1.7391748 6.184222e+07 8.200403e-02
## race.white      9.350110e+01 2.149896e+01  4.3490997 2.579984e+06 1.367028e-05
## latinx          1.148732e+00 2.345233e+01  0.0489816 3.285842e+05 9.609340e-01
## intracranialv1  3.684859e-03 7.175373e-05 51.3542419 7.511592e+07 0.000000e+00
## unmarried      -2.476365e+01 2.151760e+01 -1.1508559 1.453653e+05 2.497934e-01
## income          5.442980e-04 1.784712e-04  3.0497800 9.208567e+03 2.296599e-03
# Control Analyis in non-stigmatized group
testEstimates(sexismmod2)$estimates[, 1:5]
##                     Estimate    Std.Error    t.value          df      P(>|t|)
## (Intercept)     2.851757e+03 1.051719e+02 27.1152003 77115374.51 0.000000e+00
## sexism_factor  -1.284127e+01 1.844535e+01 -0.6961791 73331329.13 4.863167e-01
## race.white      9.870980e+01 2.211721e+01  4.4630317  3410050.57 8.081067e-06
## latinx          5.858985e+01 2.414947e+01  2.4261342   453028.51 1.526101e-02
## intracranialv1  3.412319e-03 6.665590e-05 51.1930603 73439194.95 0.000000e+00
## unmarried      -2.800381e+01 2.182705e+01 -1.2829864   226748.67 1.994982e-01
## income          7.160061e-04 1.788092e-04  4.0043022    12774.58 6.255271e-05
```

**Result: Sex/gender structural stigma was marginally associated with smaller hippocampal volume among female youth.**

H1.5a IV = Latinx ethnicity structural stigma; DVs = amygdala activation to negative vs. neutral faces, separately for right and left amygdala (tfmri_nback_all_224, tfmri_nback_all_238); covariates = age, sex at birth, marital status of household, parental education, and race; expected association = positive

```
# Multiple imputation
iamod <- immigrant_factor + race.white + sex + unmarried + income ~ 1 + (1 | site_id_l)
ia.imp <- panImpute(ss_amyg[which(ss_amyg$latinx == 1), ], formula = iamod, m = 100,
    seed = 123, silent = TRUE)
iaList <- mitmlComplete(ia.imp, print = "all")
# Primary analysis
immigrantmodla <- with(iaList, lmer(Lamyg1 ~ immigrant_factor + race.white + sex +
    unmarried + income + (1 | site_id_l/rel_family_id)))
immigrantmodra <- with(iaList, lmer(Ramyg1 ~ immigrant_factor + race.white + sex +
    unmarried + income + (1 | site_id_l/rel_family_id)))
# Control analyses in non-stigmatized youth
iamod2 <- immigrant_factor + sex + unmarried + income ~ 1 + (1 | site_id_l)
ia.imp2 <- panImpute(ss_amyg[which(ss_amyg$latinx == 0 & ss_amyg$race.white == 1),
    ], formula = iamod2, m = 100, seed = 123, silent = TRUE)
iaList2 <- mitmlComplete(ia.imp2, print = "all")
immigrantmodla2 <- with(iaList2, lmer(Lamyg1 ~ immigrant_factor + sex + unmarried +
    income + (1 | site_id_l/rel_family_id)))
immigrantmodra2 <- with(iaList2, lmer(Ramyg1 ~ immigrant_factor + sex + unmarried +
    income + (1 | site_id_l/rel_family_id)))
# Primary analysis
testEstimates(immigrantmodla)$estimates[, 1:5]
##                       Estimate    Std.Error    t.value           df   P(>|t|)
## (Intercept)       5.623803e-02 6.822321e-02  0.8243240 1.553140e+05 0.4097568
## immigrant_factor  9.043101e-03 2.950807e-02  0.3064619 6.992390e+09 0.7592530
## race.white       -2.667201e-02 4.655909e-02 -0.5728637 1.051467e+08 0.5667370
## sexM             -2.644845e-02 4.106067e-02 -0.6441309 2.045586e+10 0.5194905
## unmarried         1.694512e-02 4.631798e-02  0.3658433 3.121833e+05 0.7144822
## income            2.776333e-07 4.335185e-07  0.6404186 1.058854e+04 0.5219144
testEstimates(immigrantmodra)$estimates[, 1:5]
##                       Estimate    Std.Error    t.value           df   P(>|t|)
## (Intercept)       2.399120e-02 5.504302e-02  0.4358627 1.470284e+05 0.6629370
## immigrant_factor -1.183599e-02 2.380529e-02 -0.4972002 6.680386e+09 0.6190479
## race.white        4.303676e-02 3.754053e-02  1.1464081 1.512525e+08 0.2516263
## sexM             -4.700524e-02 3.306945e-02 -1.4214101 1.236967e+10 0.1551976
## unmarried         1.684667e-02 3.744084e-02  0.4499545 2.118443e+05 0.6527437
## income            2.604250e-07 3.490605e-07  0.7460742 1.122477e+04 0.4556382
# Control analyses in non-stigmatized youth
testEstimates(immigrantmodla2)$estimates[, 1:5]
##                       Estimate    Std.Error     t.value           df
## (Intercept)       3.773960e-02 2.341439e-02  1.61181239 2.847658e+04
## immigrant_factor -8.196777e-03 1.329208e-02 -0.61666615 2.381959e+07
## sexM              1.184503e-03 1.519097e-02  0.07797419 2.584572e+11
## unmarried        -6.329466e-02 2.153379e-02 -2.93931804 4.907185e+05
## income            1.345217e-07 1.564076e-07  0.86007147 1.531613e+04
##                      P(>|t|)
## (Intercept)      0.107013854
## immigrant_factor 0.537454955
## sexM             0.937848587
## unmarried        0.003289507
## income           0.389763107
testEstimates(immigrantmodra2)$estimates[, 1:5]
##                       Estimate    Std.Error    t.value           df    P(>|t|)
## (Intercept)       5.633634e-02 2.462535e-02  2.2877375 2.124367e+04 0.02216259
## immigrant_factor  1.440227e-02 1.465607e-02  0.9826832 1.795965e+07 0.32576338
## sexM             -2.653218e-03 1.541533e-02 -0.1721155 2.863896e+11 0.86334672
## unmarried        -6.797272e-02 2.196427e-02 -3.0946950 2.928672e+05 0.00197034
## income            1.492684e-07 1.614567e-07  0.9245106 9.949100e+03 0.35524296
```

**Result: Latinx ethnicity structural stigma was not significantly associated with amygdala reactivity among Latinx youth.**

H1.6a IV = Latinx ethnicity structural stigma; DVs = bilateral hippocampal volume, calculated by taking the mean volume across the right and left hemispheres (smri_vol_scs_hpuslh, smri_vol_scs_hpusrh); covariates = total intracranial volume, age, sex at birth, marital status of household, parental education, and race; expected association = negative

```
# Multiple imputation
ihmod <- immigrant_factor + race.white + sex + intracranialv1 + unmarried + income ~
    1 + (1 | site_id_l)
ih.imp <- panImpute(immigrant_hip, formula = ihmod, m = 100, seed = 123, silent = TRUE)
ihList <- mitmlComplete(ih.imp, print = "all")
# Primary Analysis
immigrantmod <- with(ihList, lmer(hipvol1 ~ immigrant_factor + race.white + sex +
    intracranialv1 + unmarried + income + (1 | site_id_l/rel_family_id)))
# Control Analysis in non-stigmatized group
ihmod2 <- immigrant_factor + sex + intracranialv1 + unmarried + income ~ 1 + (1 |
    site_id_l)
ih.imp2 <- panImpute(ss_hip[which(ss_hip$latinx == 0 & ss_hip$race.white == 1), ],
    formula = ihmod2, m = 100, seed = 123, silent = TRUE)
ihList2 <- mitmlComplete(ih.imp2, print = "all")
immigrantmod2 <- with(ihList2, lmer(hipvol1 ~ immigrant_factor + sex + intracranialv1 +
    unmarried + income + (1 | site_id_l/rel_family_id)))
# Sensitivity analysis within only participants who themselves or whose parents
# are Latinx immigrants
ih.imp3 <- panImpute(immigrant_hip[which(immigrant_hip$latim == 1), ], formula = ihmod,
    m = 100, seed = 123, silent = TRUE)
ihList3 <- mitmlComplete(ih.imp3, print = "all")
immigrantmod3 <- with(ihList3, lmer(hipvol1 ~ immigrant_factor + race.white + sex +
    intracranialv1 + unmarried + income + (1 | site_id_l/rel_family_id)))
# Sensitivity analysis within only participants who themselves and whose
# immediate family are not immigrants
ih.imp4 <- panImpute(immigrant_hip[which(immigrant_hip$latim == 0), ], formula = ihmod,
    m = 100, seed = 123, silent = TRUE)
ihList4 <- mitmlComplete(ih.imp4, print = "all")
immigrantmod4 <- with(ihList4, lmer(hipvol1 ~ immigrant_factor + race.white + sex +
    intracranialv1 + unmarried + income + (1 | site_id_l/rel_family_id)))
# Primary Analysis
testEstimates(immigrantmod)$estimates[, 1:5]
##                       Estimate    Std.Error     t.value           df
## (Intercept)       2.845115e+03 1.435259e+02 19.82300571 2.530488e+08
## immigrant_factor -4.010327e+01 1.990388e+01 -2.01484637 1.240691e+09
## race.white        3.215356e+01 2.838471e+01  1.13277752 4.313964e+07
## sexM              1.131748e+02 2.736924e+01  4.13510871 1.322785e+08
## intracranialv1    3.389197e-03 9.795261e-05 34.60036900 1.731061e+07
## unmarried         1.577878e+00 2.856874e+01  0.05523092 1.461285e+05
## income            5.838060e-04 2.809929e-04  2.07765393 5.489112e+03
##                       P(>|t|)
## (Intercept)      0.000000e+00
## immigrant_factor 4.392075e-02
## race.white       2.573077e-01
## sexM             3.547869e-05
## intracranialv1   0.000000e+00
## unmarried        9.559546e-01
## income           3.778762e-02
# Control Analysis in non-stigmatized group
testEstimates(immigrantmod2)$estimates[, 1:5]
##                       Estimate    Std.Error    t.value           df
## (Intercept)       2.740463e+03 9.803080e+01 27.9551232 4.786733e+07
## immigrant_factor  6.277980e+00 2.246585e+01  0.2794455 9.162483e+07
## sexM              3.895160e+01 1.737068e+01  2.2423768 1.089422e+09
## intracranialv1    3.532802e-03 6.462492e-05 54.6662506 1.774755e+08
## unmarried        -3.463533e+01 2.218353e+01 -1.5613083 3.938069e+05
## income            6.512619e-04 1.647263e-04  3.9535991 1.551765e+04
##                       P(>|t|)
## (Intercept)      0.0000000000
## immigrant_factor 0.7799029522
## sexM             0.0249370300
## intracranialv1   0.0000000000
## unmarried        0.1184518258
## income           0.0000773259
# Sensitivity analysis within only participants who themselves or whose parents
# are Latinx immigrants
testEstimates(immigrantmod3)$estimates[, 1:5]
##                      Estimate    Std.Error    t.value           df    P(>|t|)
## (Intercept)      3.070857e+03 1.951770e+02 15.7337014 1.094161e+09 0.00000000
## immigrant_factor 1.951448e+01 3.508208e+01  0.5562521 6.116861e+09 0.57803854
## race.white       5.067778e+00 3.686168e+01  0.1374809 4.228607e+07 0.89065066
## sexM             9.099115e+01 3.562954e+01  2.5538122 2.777615e+08 0.01065507
## intracranialv1   3.296816e-03 1.325257e-04 24.8768046 8.951667e+06 0.00000000
## unmarried        2.735791e+01 3.650048e+01  0.7495217 1.622703e+05 0.45354389
## income           5.441121e-04 4.114085e-04  1.3225592 5.264031e+03 0.18603947
# Sensitivity analysis within only participants who themselves and whose
# immediate family are not immigrants
testEstimates(immigrantmod4)$estimates[, 1:5]
##                       Estimate    Std.Error    t.value           df     P(>|t|)
## (Intercept)       2.503620e+03 2.272022e+02 11.0193488 1.586042e+08 0.000000000
## immigrant_factor -8.288023e+01 3.001392e+01 -2.7613930 6.257949e+08 0.005755537
## race.white        8.484961e+01 4.516100e+01  1.8788248 2.263242e+07 0.060268426
## sexM              1.245361e+02 4.298190e+01  2.8974079 1.230967e+08 0.003762603
## intracranialv1    3.573833e-03 1.547047e-04 23.1009966 3.302912e+07 0.000000000
## unmarried        -3.336900e+01 4.598101e+01 -0.7257127 1.073478e+05 0.468016511
## income            5.453932e-04 4.079085e-04  1.3370477 8.058618e+03 0.181244806
```

**Result: Latinx ethnicity structural stigma was significantly associated with smaller hippocampal volume among Latinx youth. Latinx ethnicity structural stigma was not significantly associated with hippocampal volume among Latinx youth who had immigrated themselves or whose parents were immigrants. Conversely, Latinx ethnicity structural stigma was significantly associated with smaller hippocampal volume among Latinx youth who themselves and whose parents were born in the USA.**

H1.7a IV = Race structural stigma; DVs = amygdala activation to negative vs. neutral faces, separately for right and left amygdala (tfmri_nback_all_224, tfmri_nback_all_238); covariates = age, sex at birth, marital status of household, parental education, and ethnicity; expected association = positive

```
# Multiple imputation
ramod <- racism_factor + sex + latinx + unmarried + income ~ 1 + (1 | site_id_l)
ra.imp <- panImpute(ss_amyg[which(ss_amyg$race.black == 1), ], formula = ramod, m = 100,
    seed = 123, silent = TRUE)
raList <- mitmlComplete(ra.imp, print = "all")
# Primary analysis
racismmodla <- with(raList, lmer(Lamyg1 ~ racism_factor + sex + latinx + unmarried +
    income + (1 | site_id_l/rel_family_id)))
racismmodra <- with(raList, lmer(Ramyg1 ~ racism_factor + sex + latinx + unmarried +
    income + (1 | site_id_l/rel_family_id)))
testEstimates(racismmodra)
# Control analyses in non-stigmatized group
ra.imp2 <- panImpute(ss_amyg[which(ss_amyg$race.white == "1"), ], formula = ramod,
    m = 100, seed = 123, silent = TRUE)
raList2 <- mitmlComplete(ra.imp2, print = "all")
racismmodla2 <- with(raList2, lmer(Lamyg1 ~ racism_factor + sex + latinx + unmarried +
    income + (1 | site_id_l/rel_family_id)))
racismmodra2 <- with(raList2, lmer(Ramyg1 ~ racism_factor + sex + latinx + unmarried +
    income + (1 | site_id_l/rel_family_id)))
# Primary analysis
testEstimates(racismmodla)$estimates[, 1:5]
##                    Estimate    Std.Error    t.value           df    P(>|t|)
## (Intercept)    1.294412e-01 7.401772e-02  1.7487861 1.772848e+04 0.08034531
## racism_factor  1.770060e-02 4.200398e-02  0.4214029 2.397753e+07 0.67346088
## sexM          -4.579014e-02 4.637776e-02 -0.9873297 7.267475e+08 0.32348101
## latinx         1.279145e-02 8.259423e-02  0.1548710 4.173847e+08 0.87692298
## unmarried     -2.800922e-02 6.028099e-02 -0.4646443 4.063640e+04 0.64218867
## income        -1.002774e-06 5.646549e-07 -1.7759055 6.236168e+03 0.07579724
testEstimates(racismmodra)$estimates[, 1:5]
##                    Estimate    Std.Error     t.value           df    P(>|t|)
## (Intercept)    1.306499e-01 6.495965e-02  2.01124659 1.761165e+04 0.04431462
## racism_factor  4.353170e-02 3.169243e-02  1.37356760 1.784747e+07 0.16957598
## sexM          -5.842455e-02 4.245518e-02 -1.37614644 6.253565e+09 0.16877630
## latinx        -3.794719e-02 7.425245e-02 -0.51105639 1.393234e+08 0.60931157
## unmarried      8.674426e-04 5.437190e-02  0.01595388 4.104309e+04 0.98727126
## income        -5.650079e-07 5.058055e-07 -1.11704582 6.414121e+03 0.26401658
# Control analyses in non-stigmatized group
testEstimates(racismmodla2)$estimates[, 1:5]
##                    Estimate    Std.Error    t.value           df    P(>|t|)
## (Intercept)    3.220415e-02 2.218341e-02  1.4517222 3.502679e+04 0.14658782
## racism_factor -6.937745e-03 9.574656e-03 -0.7245947 5.643864e+07 0.46870071
## sexM          -8.971272e-03 1.443915e-02 -0.6213159 1.755851e+11 0.53439178
## latinx         5.749242e-03 1.978569e-02  0.2905757 1.894208e+06 0.77137588
## unmarried     -4.047992e-02 1.955463e-02 -2.0700934 4.677833e+05 0.03844415
## income         2.016876e-07 1.473471e-07  1.3687927 1.821147e+04 0.17108098
testEstimates(racismmodra2)$estimates[, 1:5]
##                    Estimate    Std.Error    t.value           df    P(>|t|)
## (Intercept)    4.630930e-02 2.230762e-02  2.0759410 3.064330e+04 0.03790772
## racism_factor -1.048632e-02 9.624138e-03 -1.0895855 6.265955e+07 0.27589576
## sexM          -1.285328e-02 1.450084e-02 -0.8863822 8.818995e+10 0.37541163
## latinx         3.916724e-02 1.994080e-02  1.9641759 4.597654e+05 0.04951030
## unmarried     -4.352674e-02 1.960182e-02 -2.2205454 5.916643e+05 0.02638214
## income         1.843881e-07 1.480406e-07  1.2455234 1.699032e+04 0.21295659
```

**Result: Race structural stigma was not significantly associated with amygdala reactivity among Black youth.**

H1.8a IV = Race structural stigma; DVs = bilateral hippocampal volume, calculated by taking the mean volume across the right and left hemispheres (smri_vol_scs_hpuslh, smri_vol_scs_hpusrh); covariates = total intracranial volume, age, sex at birth, marital status of household, parental education, and ethnicity; expected association = negative

```
# Multiple imputation
rhmod <- racism_factor + sex + latinx + intracranialv1 + unmarried + income ~ 1 +
    (1 | site_id_l)
rh.imp <- panImpute(racism_hip, formula = rhmod, m = 100, seed = 123, silent = TRUE)
rhList <- mitmlComplete(rh.imp, print = "all")
# Primary analysis
racismmod <- with(rhList, lmer(hipvol1 ~ racism_factor + sex + latinx + intracranialv1 +
    unmarried + income + (1 | site_id_l/rel_family_id)))
# Control analysis in non-stigmatized group
rh.imp2 <- panImpute(ss_hip[which(ss_hip$race.white == "1"), ], formula = rhmod,
    m = 100, seed = 123, silent = TRUE)
rhList2 <- mitmlComplete(rh.imp2, print = "all")
racismmod2 <- with(rhList2, lmer(hipvol1 ~ racism_factor + sex + latinx + intracranialv1 +
    unmarried + income + (1 | site_id_l/rel_family_id)))
# Primary analysis
testEstimates(racismmod)$estimates[, 1:5]
##                     Estimate    Std.Error     t.value           df    P(>|t|)
## (Intercept)     2.411495e+03 1.452708e+02 16.59999102 3.295865e+07 0.00000000
## racism_factor  -5.825933e+01 2.570025e+01 -2.26687735 2.137877e+07 0.02339773
## sexM            2.810785e+01 2.756712e+01  1.01961497 6.564279e+08 0.30791110
## latinx          8.317158e+01 4.652157e+01  1.78780700 1.444861e+06 0.07380736
## intracranialv1  3.657762e-03 1.004476e-04 36.41462402 4.056999e+07 0.00000000
## unmarried       8.283800e-01 3.303836e+01  0.02507328 5.287449e+04 0.97999661
## income          2.316192e-04 3.169590e-04  0.73075444 9.970765e+03 0.46494633
# Control analysis in non-stigmatized group
testEstimates(racismmod2)$estimates[, 1:5]
##                     Estimate    Std.Error   t.value           df      P(>|t|)
## (Intercept)     2.793145e+03 8.599805e+01 32.479161  26710200.86 0.000000e+00
## racism_factor  -2.156210e+01 1.800277e+01 -1.197710 425893893.20 2.310299e-01
## sexM            5.826927e+01 1.532831e+01  3.801415 543104684.67 1.438721e-04
## latinx          1.440033e+01 2.101527e+01  0.685232    347779.17 4.931980e-01
## intracranialv1  3.487941e-03 5.709736e-05 61.087593  92439616.82 0.000000e+00
## unmarried      -2.645538e+01 1.869802e+01 -1.414876    359393.41 1.571057e-01
## income          6.347267e-04 1.452779e-04  4.369053     15962.33 1.255738e-05
```

**Result: Race structural stigma was significantly associated with smaller hippocampal volume among Black youth.**

### Analyses involving Structural Stigma

Multiple Imputation

```
# Sexual orientation
lhmod <- sex + race.white + latinx + intracranialv1 + unmarried + income ~ 1 + (1 |
    site_id_l)
lh.imp <- panImpute(lgbc_hip1, formula = lhmod, m = 100, seed = 123, silent = TRUE)
lhList <- mitmlComplete(lh.imp, print = "all")
# Sex
samod <- race.white + latinx + unmarried + income ~ 1 + (1 | site_id_l)
sa.imp <- panImpute(ss_amyg[which(ss_amyg$sex == "F"), ], formula = samod, m = 100,
    seed = 123, silent = TRUE)
saList <- mitmlComplete(sa.imp, print = "all")

shmod <- race.white + latinx + intracranialv1 + unmarried + income ~ 1 + (1 | site_id_l)
sh.imp <- panImpute(sexism_hip, formula = shmod, m = 100, seed = 123, silent = TRUE)
shList <- mitmlComplete(sh.imp, print = "all")

# Latinx
iamod <- race.white + sex + unmarried + income ~ 1 + (1 | site_id_l)
ia.imp <- panImpute(ss_amyg[which(ss_amyg$latinx == 1), ], formula = iamod, m = 100,
    seed = 123, silent = TRUE)
iaList <- mitmlComplete(ia.imp, print = "all")

ihmod <- immigrant_factor + race.white + sex + intracranialv1 + unmarried + income ~
    1 + (1 | site_id_l)
ih.imp <- panImpute(immigrant_hip, formula = ihmod, m = 100, seed = 123, silent = TRUE)
ihList <- mitmlComplete(ih.imp, print = "all")

# Race
ramod <- racism_factor + sex + latinx + unmarried + income ~ 1 + (1 | site_id_l)
ra.imp <- panImpute(ss_amyg[which(ss_amyg$race.black == 1), ], formula = ramod, m = 100,
    seed = 123, silent = TRUE)
raList <- mitmlComplete(ra.imp, print = "all")

rhmod <- racism_factor + sex + latinx + intracranialv1 + unmarried + income ~ 1 +
    (1 | site_id_l)
rh.imp <- panImpute(racism_hip, formula = rhmod, m = 100, seed = 123, silent = TRUE)
rhList <- mitmlComplete(rh.imp, print = "all")
```

H.1.9a-12a IV = Perceived discrimination; DVs = amygdala activation to negative vs. neutral faces, separately for right and left amygdala (tfmri_nback_all_224, tfmri_nback_all_238); covariates = age, sex at birth (in models not focused on sex/gender), marital status of household, parental education, race, and ethnicity (in models not focused on race/ethnicity); expected association = positive

```
# LGB Discrimination Continuous measure
lgb_discrim_modla <- gamm4(Lamyg1 ~ discrim_cont + sex + race.white + unmarried +
    income, random = ~(1 | site_id_l), data = lgbc_amyg1)
lgb_discrim_modra <- gamm4(Ramyg1 ~ discrim_cont + sex + race.white + unmarried +
    income, random = ~(1 | site_id_l), data = lgbc_amyg1)
# Dichotomous measure
lgb_discrim_modla2 <- gamm4(Lamyg1 ~ dim_yesno_q3 + sex + race.white + unmarried +
    income, random = ~(1 | site_id_l), data = lgbc_amyg1)
lgb_discrim_modra2 <- gamm4(Ramyg1 ~ dim_yesno_q3 + sex + race.white + unmarried +
    income, random = ~(1 | site_id_l), data = lgbc_amyg1)
# Sex-based discrimination
sex_disrim_modla <- with(saList, lmer(Lamyg1 ~ discrim_cont + race.white + latinx +
    unmarried + income + (1 | site_id_l/rel_family_id)))
sex_disrim_modra <- with(saList, lmer(Ramyg1 ~ discrim_cont + race.white + latinx +
    unmarried + income + (1 | site_id_l/rel_family_id)))
# Latinx discrimination Continuous measure
latinx_discrim_modla <- with(iaList, lmer(Lamyg1 ~ latinx_discrim + race.white +
    sex + unmarried + income + (1 | site_id_l/rel_family_id)))
latinx_discrim_modra <- with(iaList, lmer(Ramyg1 ~ latinx_discrim + race.white +
    sex + unmarried + income + (1 | site_id_l/rel_family_id)))
# Dichotomous measure
latinx_discrim_modla2 <- with(iaList, lmer(Lamyg1 ~ latinx_discrim_dic + race.white +
    sex + unmarried + income + (1 | site_id_l/rel_family_id)))
latinx_discrim_modra2 <- with(iaList, lmer(Ramyg1 ~ latinx_discrim_dic + race.white +
    sex + unmarried + income + (1 | site_id_l/rel_family_id)))
# Race discrimination Continuous measure
race_discrim_modla <- with(raList, lmer(Lamyg1 ~ discrim_cont + sex + latinx + unmarried +
    income + (1 | site_id_l/rel_family_id)))
race_discrim_modra <- with(raList, lmer(Ramyg1 ~ discrim_cont + sex + latinx + unmarried +
    income + (1 | site_id_l/rel_family_id)))
# Dichotomous measure
race_discrim_modla2 <- with(raList, lmer(Lamyg1 ~ dim_yesno_q1 + sex + latinx + unmarried +
    income + (1 | site_id_l/rel_family_id)))
race_discrim_modra2 <- with(raList, lmer(Ramyg1 ~ dim_yesno_q1 + sex + latinx + unmarried +
    income + (1 | site_id_l/rel_family_id)))
# LGB Discrimination Continuous measure
summary(lgb_discrim_modla$gam)
##
## Family: gaussian
## Link function: identity
##
## Formula:
## Lamyg1 ~ discrim_cont + sex + race.white + unmarried + income
##
## Parametric coefficients:
##                Estimate Std. Error t value Pr(>|t|)
## (Intercept)  -6.544e-01  3.669e-01  -1.784   0.0862 .
## discrim_cont  2.673e-01  1.466e-01   1.823   0.0798 .
## sexM          8.009e-02  1.333e-01   0.601   0.5531
## race.white    2.241e-01  2.092e-01   1.071   0.2940
## unmarried     2.043e-02  2.024e-01   0.101   0.9204
## income        9.689e-07  1.338e-06   0.724   0.4756
## ---
## Signif. codes:  0 '***' 0.001 '**' 0.01 '*' 0.05 '.' 0.1 ' ' 1
##
##
## R-sq.(adj) =  -0.0798
## lmer.REML = 60.501  Scale est. = 0.094649  n = 32
summary(lgb_discrim_modra$gam)
##
## Family: gaussian
## Link function: identity
##
## Formula:
## Ramyg1 ~ discrim_cont + sex + race.white + unmarried + income
##
## Parametric coefficients:
##                Estimate Std. Error t value Pr(>|t|)
## (Intercept)  -9.912e-01  6.672e-01  -1.486    0.149
## discrim_cont  1.397e-01  2.765e-01   0.505    0.618
## sexM          1.981e-01  2.465e-01   0.804    0.429
## race.white    6.229e-01  3.655e-01   1.704    0.100
## unmarried     5.160e-01  3.505e-01   1.472    0.153
## income        2.925e-06  2.485e-06   1.177    0.250
##
##
## R-sq.(adj) =  0.00735
## lmer.REML = 88.025  Scale est. = 0.43951   n = 32
# Dichotomous measure
summary(lgb_discrim_modla2$gam)
##
## Family: gaussian
## Link function: identity
##
## Formula:
## Lamyg1 ~ dim_yesno_q3 + sex + race.white + unmarried + income
##
## Parametric coefficients:
##                Estimate Std. Error t value Pr(>|t|)
## (Intercept)  -2.822e-01  3.250e-01  -0.868    0.394
## dim_yesno_q3  6.953e-02  1.865e-01   0.373    0.713
## sexM         -1.801e-02  1.529e-01  -0.118    0.907
## race.white    9.986e-02  2.143e-01   0.466    0.645
## unmarried     3.102e-02  2.240e-01   0.138    0.891
## income        1.191e-06  1.764e-06   0.675    0.506
##
##
## R-sq.(adj) =  -0.169
## lmer.REML = 61.674  Scale est. = 0.11756   n = 30
summary(lgb_discrim_modra2$gam)
##
## Family: gaussian
## Link function: identity
##
## Formula:
## Ramyg1 ~ dim_yesno_q3 + sex + race.white + unmarried + income
##
## Parametric coefficients:
##                Estimate Std. Error t value Pr(>|t|)
## (Intercept)  -4.853e-01  5.332e-01  -0.910    0.372
## dim_yesno_q3 -2.966e-01  3.168e-01  -0.936    0.358
## sexM          1.417e-01  2.697e-01   0.525    0.604
## race.white    5.692e-01  3.592e-01   1.585    0.126
## unmarried     4.510e-01  3.644e-01   1.238    0.228
## income        1.258e-06  2.954e-06   0.426    0.674
##
##
## R-sq.(adj) =  0.012
## lmer.REML = 84.691  Scale est. = 0.46325   n = 30
# Sex-based discrimination
testEstimates(sex_disrim_modla)$estimates[, 1:5]
##                   Estimate    Std.Error    t.value          df   P(>|t|)
## (Intercept)   2.844812e-02 4.993941e-02  0.5696526    556126.4 0.5689135
## discrim_cont  9.165953e-03 3.008973e-02  0.3046206 692922883.8 0.7606551
## race.white   -2.979713e-02 2.741985e-02 -1.0866992  16402671.4 0.2771698
## latinx        1.666606e-02 2.765837e-02  0.6025682   2137804.6 0.5467961
## unmarried     2.471329e-02 2.766618e-02  0.8932672    781146.7 0.3717144
## income        3.160476e-07 2.164296e-07  1.4602791     35740.1 0.1442222
testEstimates(sex_disrim_modra)$estimates[, 1:5]
##                   Estimate    Std.Error     t.value           df   P(>|t|)
## (Intercept)   5.156743e-02 4.658446e-02  1.10696638    747619.04 0.2683088
## discrim_cont -2.259067e-03 2.811189e-02 -0.08035984 917855684.10 0.9359511
## race.white   -1.393285e-02 2.558277e-02 -0.54461865  24016089.33 0.5860158
## latinx        2.484032e-02 2.589810e-02  0.95915615    589431.33 0.3374805
## unmarried     1.158075e-02 2.579106e-02  0.44902173   1154825.56 0.6534161
## income        2.137160e-07 2.011377e-07  1.06253564     49252.36 0.2879978
# Latinx discrimination Continuous measure
testEstimates(latinx_discrim_modla)$estimates[, 1:5]
##                     Estimate    Std.Error    t.value           df   P(>|t|)
## (Intercept)     4.135719e-02 7.821700e-02  0.5287493 2.751417e+05 0.5969798
## latinx_discrim  1.016957e-02 4.158483e-02  0.2445501 2.977823e+08 0.8068048
## race.white     -2.346782e-02 4.543589e-02 -0.5165039 8.286110e+07 0.6055025
## sexM           -2.732486e-02 4.115896e-02 -0.6638861 3.142687e+10 0.5067632
## unmarried       4.869333e-03 4.622136e-02  0.1053481 4.084263e+05 0.9160997
## income          2.314146e-07 4.319770e-07  0.5357103 1.416712e+04 0.5921672
testEstimates(latinx_discrim_modra)$estimates[, 1:5]
##                     Estimate    Std.Error     t.value           df   P(>|t|)
## (Intercept)     4.254632e-02 6.145168e-02  0.69235404 9.695133e+04 0.4887167
## latinx_discrim  4.779662e-03 3.243954e-02  0.14734061 1.037848e+08 0.8828632
## race.white      4.384747e-02 3.548743e-02  1.23557747 3.337522e+07 0.2166157
## sexM           -4.025854e-02 3.210984e-02 -1.25377580 2.302704e+10 0.2099235
## unmarried       2.147801e-03 3.624850e-02  0.05925215 1.786385e+05 0.9527513
## income          1.472464e-07 3.470010e-07  0.42434007 5.497399e+03 0.6713344
# Dichotomous measure
testEstimates(latinx_discrim_modla2)$estimates[, 1:5]
##                         Estimate    Std.Error    t.value           df
## (Intercept)         3.399789e-02 5.553296e-02  0.6122110 8.844489e+04
## latinx_discrim_dic  1.262315e-01 7.465688e-02  1.6908220 1.286355e+08
## race.white         -7.163699e-03 4.269918e-02 -0.1677714 6.550276e+07
## sexM               -4.804908e-02 3.868022e-02 -1.2422133 1.095162e+10
## unmarried           3.642355e-02 4.367127e-02  0.8340392 2.724356e+05
## income              2.753072e-07 4.103986e-07  0.6708289 1.027964e+04
##                       P(>|t|)
## (Intercept)        0.54039975
## latinx_discrim_dic 0.09087081
## race.white         0.86676316
## sexM               0.21415789
## unmarried          0.40425962
## income             0.50234463
testEstimates(latinx_discrim_modra2)$estimates[, 1:5]
##                         Estimate    Std.Error     t.value           df
## (Intercept)         4.329663e-02 4.826357e-02  0.89708717 4.283790e+04
## latinx_discrim_dic -1.030441e-03 6.400082e-02 -0.01610044 6.942508e+07
## race.white          3.541504e-02 3.682999e-02  0.96158158 3.582701e+07
## sexM               -6.249823e-02 3.321514e-02 -1.88161864 4.399383e+09
## unmarried           1.702080e-02 3.783666e-02  0.44984954 1.581780e+05
## income              3.125845e-07 3.616345e-07  0.86436566 5.322866e+03
##                       P(>|t|)
## (Intercept)        0.36967745
## latinx_discrim_dic 0.98715426
## race.white         0.33625984
## sexM               0.05988781
## unmarried          0.65281955
## income             0.38742600
# Race discrimination Continuous measure
testEstimates(race_discrim_modla)$estimates[, 1:5]
##                   Estimate    Std.Error    t.value           df    P(>|t|)
## (Intercept)   7.701684e-02 9.172215e-02  0.8396755 2.592674e+04 0.40109810
## discrim_cont  5.392812e-02 4.252563e-02  1.2681323 8.065175e+08 0.20475072
## sexM         -7.431873e-02 4.689847e-02 -1.5846729 7.724217e+08 0.11304066
## latinx        2.001973e-02 8.401382e-02  0.2382909 3.889337e+08 0.81165550
## unmarried    -2.943811e-02 6.256268e-02 -0.4705379 2.869838e+04 0.63797431
## income       -9.630759e-07 5.826858e-07 -1.6528219 4.941022e+03 0.09843065
testEstimates(race_discrim_modra)$estimates[, 1:5]
##                   Estimate    Std.Error      t.value           df   P(>|t|)
## (Intercept)   8.512363e-02 8.176462e-02  1.041081526 3.909795e+04 0.2978442
## discrim_cont  5.313031e-02 4.091720e-02  1.298483451 2.079735e+09 0.1941213
## sexM         -5.554206e-02 4.377540e-02 -1.268796117 2.605992e+09 0.2045138
## latinx       -4.962723e-02 7.537658e-02 -0.658390604 1.100447e+08 0.5102872
## unmarried    -1.592396e-04 5.586702e-02 -0.002850334 4.184050e+04 0.9977258
## income       -7.296447e-07 5.143408e-07 -1.418601605 6.450319e+03 0.1560635
# Dichotomous measure
testEstimates(race_discrim_modla2)$estimates[, 1:5]
##                   Estimate    Std.Error     t.value           df    P(>|t|)
## (Intercept)   9.334982e-02 6.181998e-02  1.51002653 9.797636e+03 0.13106890
## dim_yesno_q1  8.418461e-02 6.715690e-02  1.25355111 9.984747e+08 0.21000521
## sexM         -1.344843e-02 3.930537e-02 -0.34215260 5.829580e+08 0.73223606
## latinx        6.537922e-03 6.931998e-02  0.09431511 8.503621e+07 0.92485885
## unmarried    -2.597517e-02 5.135752e-02 -0.50577154 2.280007e+04 0.61302193
## income       -9.652365e-07 4.851933e-07 -1.98938531 3.481133e+03 0.04673682
testEstimates(race_discrim_modra2)$estimates[, 1:5]
```

```
##                   Estimate    Std.Error    t.value           df   P(>|t|)
## (Intercept)   6.761214e-02 5.767020e-02  1.1723930 9.157204e+03 0.2410698
## dim_yesno_q1  9.073540e-02 6.272182e-02  1.4466322 3.694282e+09 0.1480000
## sexM         -2.819038e-02 3.664461e-02 -0.7692915 9.310198e+09 0.4417203
## latinx       -4.212635e-02 6.457515e-02 -0.6523616 4.473285e+07 0.5141679
## unmarried     3.096245e-02 4.794433e-02  0.6458000 1.934674e+04 0.5184165
## income       -3.739792e-07 4.521198e-07 -0.8271685 3.393124e+03 0.4081997
```

**Result: Perceived discrimination (when measured dichotomously) was significantly associated with greater left amygdala reactivity to negative vs. neutral faces in Latinx youth. No other significant associations were observed between perceived discrimination and amygdala reactivity.**

H.1.13a-16a IV = Perceived discrimination (1 year follow-up). DVs = bilateral hippocampal volume (baseline), calculated by taking the mean volume across the right and left hemispheres (smri_vol_scs_hpuslh, smri_vol_scs_hpusrh); covariates = age, sex at birth (in models not focused on sex/gender), marital status of household, parental education, race, and ethnicity (in models not focused on race/ethnicity); expected association = negative

```
# LGB Discrimination Continuous measure
lgb_discrim_mod <- with(lhList, lmer(hipvol1 ~ discrim_cont + sex + race.white +
    latinx + intracranialv1 + unmarried + income + (1 | site_id_l)))
# Dichotomous measure
lgb_discrim_mod2 <- with(lhList, lmer(hipvol1 ~ dim_yesno_q3 + sex + race.white +
    latinx + intracranialv1 + unmarried + income + (1 | site_id_l)))
# Sex-based discrimination
sex_disrim_mod <- with(shList, lmer(hipvol1 ~ discrim_cont + race.white + latinx +
    intracranialv1 + unmarried + income + (1 | site_id_l/rel_family_id)))
# Latinx discrimination Continuous measure
latinx_discrim_mod <- with(ihList, lmer(hipvol1 ~ latinx_discrim + race.white + sex +
    intracranialv1 + unmarried + income + (1 | site_id_l/rel_family_id)))
# Dichotomous measure
latinx_discrim_mod2 <- with(ihList, lmer(hipvol1 ~ latinx_discrim_dic + race.white +
    sex + intracranialv1 + unmarried + income + (1 | site_id_l/rel_family_id)))
# Race discrimination Continuous measure
race_discrim_mod <- with(rhList, lmer(hipvol1 ~ discrim_cont + sex + latinx + intracranialv1 +
    unmarried + income + (1 | site_id_l/rel_family_id)))
# Dichotomous measure
race_discrim_mod2 <- with(rhList, lmer(hipvol1 ~ dim_yesno_q1 + sex + latinx + intracranialv1 +
    unmarried + income + (1 | site_id_l/rel_family_id)))
# LGB Discrimination Continuous measure
testEstimates(lgb_discrim_mod)$estimates[, 1:5]
##                     Estimate    Std.Error    t.value           df      P(>|t|)
## (Intercept)    -1.111331e+03 8.920121e+02 -1.2458696  41548123.72 0.2128122745
## discrim_cont    1.003692e+02 1.489250e+02  0.6739582 655381702.06 0.5003378627
## sexM           -5.108029e+02 1.802716e+02 -2.8335189   2583172.68 0.0046038942
## race.white      6.811168e+02 2.042240e+02  3.3351456 106788821.63 0.0008525477
## latinx          4.162154e+02 2.373993e+02  1.7532294   3533597.71 0.0795627313
## intracranialv1  5.595258e-03 6.150971e-04  9.0965451    763121.75 0.0000000000
## unmarried       2.278121e+02 2.194349e+02  1.0381765     76354.62 0.2991911804
## income         -2.096865e-04 1.699326e-03 -0.1233939     10817.57 0.9017974420
# Dichotomous measure
testEstimates(lgb_discrim_mod2)$estimates[, 1:5]
##                     Estimate    Std.Error    t.value          df      P(>|t|)
## (Intercept)    -1.276393e+03 9.201815e+02 -1.3871095 5370101.226 0.1654084199
## dim_yesno_q3   -9.993937e+01 1.906807e+02 -0.5241190  152002.659 0.6001965643
## sexM           -5.354455e+02 1.813805e+02 -2.9520566 1581766.299 0.0031566971
## race.white      7.127008e+02 2.080026e+02  3.4264032 3742798.368 0.0006116383
## latinx          4.084343e+02 2.352433e+02  1.7362208 4175575.523 0.0825248730
## intracranialv1  5.863246e-03 6.732097e-04  8.7093896  295413.218 0.0000000000
## unmarried       1.739094e+02 2.238324e+02  0.7769628   64540.947 0.4371835602
## income         -1.011686e-03 2.010988e-03 -0.5030790    8229.581 0.6149221822
# Sex-based discrimination
testEstimates(sex_disrim_mod)$estimates[, 1:5]
##                     Estimate    Std.Error    t.value           df      P(>|t|)
## (Intercept)     2.449166e+03 1.119647e+02 21.8744550  60387752.35 0.000000e+00
## discrim_cont   -1.312906e+01 2.147814e+01 -0.6112752 110346453.48 5.410174e-01
## race.white      1.044732e+02 2.258864e+01  4.6250318   4060213.90 3.745524e-06
## latinx          1.139049e+01 2.442528e+01  0.4663400    375797.55 6.409724e-01
## intracranialv1  3.665988e-03 7.448565e-05 49.2173719  52563632.75 0.000000e+00
## unmarried      -1.341920e+01 2.249110e+01 -0.5966448    142554.80 5.507455e-01
## income          6.076442e-04 1.841404e-04  3.2998952     10779.82 9.703484e-04
# Latinx discrimination Continuous measure
testEstimates(latinx_discrim_mod)$estimates[, 1:5]
##                     Estimate    Std.Error    t.value           df      P(>|t|)
## (Intercept)     3.017140e+03 1.501081e+02 20.0997830 2.056037e+08 0.0000000000
## latinx_discrim -4.189047e+01 2.554329e+01 -1.6399793 7.076360e+07 0.1010094689
## race.white      3.784798e+01 2.966941e+01  1.2756565 3.078793e+07 0.2020769940
## sexM            1.237119e+02 2.871387e+01  4.3084361 3.395289e+08 0.0000164413
## intracranialv1  3.331016e-03 1.031149e-04 32.3039229 1.790699e+07 0.0000000000
## unmarried       6.388327e+00 2.999958e+01  0.2129472 1.440084e+05 0.8313684269
## income          4.364507e-04 2.951658e-04  1.4786628 5.382484e+03 0.1392889869
# Dichotomous measure
testEstimates(latinx_discrim_mod2)$estimates[, 1:5]
##                         Estimate    Std.Error     t.value           df
## (Intercept)         2.942257e+03 1.500172e+02 19.61279946 8.449451e+08
## latinx_discrim_dic -1.081202e+01 4.843054e+01 -0.22324804 2.732053e+08
## race.white          4.392410e+01 3.018193e+01  1.45531103 2.427400e+07
## sexM                1.196544e+02 2.915707e+01  4.10378683 2.557738e+08
## intracranialv1      3.353260e-03 1.055248e-04 31.77699579 1.118895e+07
## unmarried          -1.198066e+00 3.056592e+01 -0.03919614 1.343139e+05
## income              3.808770e-04 3.008299e-04  1.26608776 5.105977e+03
##                         P(>|t|)
## (Intercept)        0.000000e+00
## latinx_discrim_dic 8.233425e-01
## race.white         1.455832e-01
## sexM               4.064421e-05
## intracranialv1     0.000000e+00
## unmarried          9.687341e-01
## income             2.055394e-01
# Race discrimination Continuous measure
testEstimates(race_discrim_mod)$estimates[, 1:5]
##                    Estimate    Std.Error    t.value           df    P(>|t|)
## (Intercept)    2.398626e+03 1.562239e+02 15.3537694 1.633535e+07 0.00000000
## discrim_cont   1.596828e+01 2.248132e+01  0.7102910 9.662695e+08 0.47752372
## sexM           4.588768e+01 2.953971e+01  1.5534237 9.578415e+08 0.12032193
## latinx         9.526024e+01 4.925065e+01  1.9341924 2.218410e+06 0.05308961
## intracranialv1 3.628754e-03 1.053326e-04 34.4504361 4.106700e+07 0.00000000
## unmarried      2.192200e+01 3.569083e+01  0.6142194 3.570538e+04 0.53907426
## income         3.538659e-04 3.368132e-04  1.0506295 7.201908e+03 0.29346404
# Dichotomous measure
testEstimates(race_discrim_mod2)$estimates[, 1:5]
##                     Estimate    Std.Error    t.value           df   P(>|t|)
## (Intercept)     2.361199e+03 1.603221e+02 14.7278377 1.442504e+07 0.0000000
## dim_yesno_q1   -1.017900e+01 4.396598e+01 -0.2315200 2.957843e+10 0.8169108
## sexM            4.165585e+01 3.088706e+01  1.3486506 6.760223e+08 0.1774492
## latinx          9.183516e+01 5.209501e+01  1.7628399 2.926834e+06 0.0779276
## intracranialv1  3.691662e-03 1.105849e-04 33.3830718 4.716585e+07 0.0000000
## unmarried      -8.033248e+00 3.737890e+01 -0.2149140 2.889006e+04 0.8298359
## income          1.059198e-04 3.585515e-04  0.2954104 6.054978e+03 0.7676905
```

**Result: No significant associations between perceived discrimination and hippocampal volume were found.**

H.1.13a-16a IV = Perceived discrimination (1 year follow-up). DVs = bilateral hippocampal volume (2-year follow-up)

```
# LGB Discrimination Continuous measure
lgb_discrim_mod <- with(lhList, lmer(hipvol2 ~ discrim_cont + sex + race.white +
    latinx + intracranialv1 + unmarried + income + (1 | site_id_l)))
# Dichotomous measure
lgb_discrim_mod2 <- with(lhList, lmer(hipvol2 ~ dim_yesno_q3 + sex + race.white +
    latinx + intracranialv1 + unmarried + income + (1 | site_id_l)))
# Sex-based discrimination
sex_disrim_mod <- with(shList, lmer(hipvol2 ~ discrim_cont + race.white + latinx +
    intracranialv1 + unmarried + income + (1 | site_id_l/rel_family_id)))
# Latinx discrimination Continuous measure
latinx_discrim_mod <- with(ihList, lmer(hipvol2 ~ latinx_discrim + race.white + sex +
    intracranialv1 + unmarried + income + (1 | site_id_l/rel_family_id)))
# Dichotomous measure
latinx_discrim_mod2 <- with(ihList, lmer(hipvol2 ~ latinx_discrim_dic + race.white +
    sex + intracranialv1 + unmarried + income + (1 | site_id_l/rel_family_id)))
# Race discrimination Continuous measure
race_discrim_mod <- with(rhList, lmer(hipvol2 ~ discrim_cont + sex + latinx + intracranialv1 +
    unmarried + income + (1 | site_id_l/rel_family_id)))
# Dichotomous measure
race_discrim_mod2 <- with(rhList, lmer(hipvol2 ~ dim_yesno_q1 + sex + latinx + intracranialv1 +
    unmarried + income + (1 | site_id_l/rel_family_id)))
# LGB Discrimination Continuous measure
testEstimates(lgb_discrim_mod)$estimates[, 1:5]
##                     Estimate    Std.Error     t.value           df      P(>|t|)
## (Intercept)    -1.621344e+03 1.724508e+03 -0.94017774   1856653.84 3.471265e-01
## discrim_cont   -3.719043e+02 4.055559e+02 -0.91702354  23963347.44 3.591303e-01
## sexM           -5.688641e+02 2.947675e+02 -1.92987360 165131809.95 5.362250e-02
## race.white      7.808993e+02 3.579946e+02  2.18131575  16627667.75 2.916008e-02
## latinx          6.212845e+02 4.017829e+02  1.54631903   5473713.94 1.220276e-01
## intracranialv1  6.266318e-03 1.224443e-03  5.11769073    234055.36 3.095430e-07
## unmarried       2.323074e+02 3.370793e+02  0.68917739    146727.25 4.907127e-01
## income          1.454009e-04 2.550675e-03  0.05700487     12949.61 9.545422e-01
# Dichotomous measure
testEstimates(lgb_discrim_mod2)$estimates[, 1:5]
##                     Estimate    Std.Error      t.value           df
## (Intercept)    -1.590935e+03 1.752702e+03 -0.907704250   2122772.55
## dim_yesno_q3   -1.941121e+02 2.818895e+02 -0.688610499   1834056.60
## sexM           -6.582339e+02 3.120379e+02 -2.109467417  19769018.47
## race.white      8.152030e+02 3.924084e+02  2.077435421 147793527.59
## latinx          6.743569e+02 4.071214e+02  1.656402565   5588760.89
## intracranialv1  6.037888e-03 1.192618e-03  5.062718290    320482.95
## unmarried       1.989578e+02 3.362179e+02  0.591752565    224898.74
## income         -2.552455e-05 2.658485e-03 -0.009601166     16432.33
##                     P(>|t|)
## (Intercept)    3.640346e-01
## dim_yesno_q3   4.910685e-01
## sexM           3.490427e-02
## race.white     3.776139e-02
## latinx         9.764038e-02
## intracranialv1 4.135485e-07
## unmarried      5.540169e-01
## income         9.923396e-01
# Sex-based discrimination
testEstimates(sex_disrim_mod)$estimates[, 1:5]
##                     Estimate    Std.Error   t.value           df    P(>|t|)
## (Intercept)     2.844729e+03 1.612185e+02 17.645179  51156296.83 0.00000000
## discrim_cont   -4.774052e+01 3.244387e+01 -1.471480 387675360.94 0.14116127
## race.white      7.291561e+01 3.341163e+01  2.182343   7026365.04 0.02908429
## latinx         -4.602317e+01 3.530154e+01 -1.303715    338447.43 0.19233149
## intracranialv1  3.537220e-03 1.073524e-04 32.949616 214870864.52 0.00000000
## unmarried      -3.767786e+01 3.262193e+01 -1.154986    346104.68 0.24809713
## income          5.837804e-04 2.691445e-04  2.169022     13782.27 0.03009802
# Latinx discrimination Continuous measure
testEstimates(latinx_discrim_mod)$estimates[, 1:5]
##                     Estimate    Std.Error    t.value           df     P(>|t|)
## (Intercept)     3.227636e+03 2.181933e+02 14.7925517  96050262.84 0.000000000
## latinx_discrim -2.566592e+01 4.119691e+01 -0.6230058  34554185.61 0.533280703
## race.white     -4.484339e+01 4.474226e+01 -1.0022603  28278696.07 0.316217895
## sexM            1.508464e+02 4.224178e+01  3.5710250 255438161.97 0.000355587
## intracranialv1  3.282370e-03 1.492260e-04 21.9959746  13892809.69 0.000000000
## unmarried      -4.452771e+01 4.513112e+01 -0.9866297    126876.91 0.323826099
## income          6.841779e-04 4.529593e-04  1.5104621      6029.66 0.130977947
# Dichotomous measure
testEstimates(latinx_discrim_mod2)$estimates[, 1:5]
##                         Estimate    Std.Error     t.value           df
## (Intercept)         3.131468e+03 2.208169e+02 14.18128541 3.296158e+08
## latinx_discrim_dic -4.698740e+00 7.151656e+01 -0.06570142 2.682706e+08
## race.white         -4.036527e+01 4.579558e+01 -0.88142275 2.497088e+07
## sexM                1.460699e+02 4.304272e+01  3.39360247 2.425010e+08
## intracranialv1      3.327777e-03 1.551917e-04 21.44301604 1.139657e+07
## unmarried          -4.431003e+01 4.620595e+01 -0.95896797 1.214415e+05
## income              6.572982e-04 4.609201e-04  1.42605695 6.107398e+03
##                         P(>|t|)
## (Intercept)        0.0000000000
## latinx_discrim_dic 0.9476155387
## race.white         0.3780890590
## sexM               0.0006897976
## intracranialv1     0.0000000000
## unmarried          0.3375767925
## income             0.1539030148
# Race discrimination Continuous measure
testEstimates(race_discrim_mod)$estimates[, 1:5]
##                     Estimate    Std.Error     t.value           df   P(>|t|)
## (Intercept)     2.347971e+03 2.365451e+02  9.92610175 4.512801e+07 0.0000000
## discrim_cont    5.414323e+01 3.407763e+01  1.58882052 4.453093e+09 0.1121009
## sexM            1.071912e+01 4.625575e+01  0.23173604 1.540180e+08 0.8167430
## latinx         -2.530026e+01 7.113073e+01 -0.35568676 6.808041e+06 0.7220752
## intracranialv1  3.734380e-03 1.605667e-04 23.25750035 3.993120e+07 0.0000000
## unmarried       3.173331e+00 5.260837e+01  0.06031988 9.712391e+04 0.9519010
## income          3.365743e-04 5.255563e-04  0.64041532 8.039864e+03 0.5219209
# Dichotomous measure
testEstimates(race_discrim_mod2)$estimates[, 1:5]
##                     Estimate    Std.Error     t.value           df   P(>|t|)
## (Intercept)     2.385145e+03 2.434322e+02  9.79798564 5.059527e+07 0.0000000
## dim_yesno_q1   -9.755402e+00 6.552489e+01 -0.14888086 5.362549e+08 0.8816476
## sexM            5.048658e+00 4.858608e+01  0.10391163 1.826306e+08 0.9172395
## latinx         -1.408945e+01 7.559291e+01 -0.18638583 2.749241e+06 0.8521422
## intracranialv1  3.757958e-03 1.691052e-04 22.22260613 3.913226e+07 0.0000000
## unmarried      -5.223018e+00 5.509867e+01 -0.09479391 9.858236e+04 0.9244787
## income          2.721602e-04 5.588146e-04  0.48703132 8.299642e+03 0.6262490
```

**Result: No significant associations between perceived discrimination and hippocampal volume were found.**

### Figures

\# Sex structural stigma
#Create ordered categorical variables of states in order of structural stigma
sexism_hip$state<-as.factor(sexism_hip$sexism_factor)
statelev<-NULL
ss_hip$STATE<-as.factor(ss_hip$STATE)
for(x in 1:17){
 statelev[x]<-ss_hip$STATE[which(as.factor(ss_hip$sexism_factor)==levels(sexism_hip$state)[x])][1]
 }
levels(sexism_hip$state)<-levels(ss_hip$STATE)[statelev]

#Graph violin plots
sexism_hip.graph<-ggplot(aes(x=state,y=hipvol1,fill=state),data=sexism_hip)+geom_violin()+theme_classic()+theme(text = element_text(size=20),axis.text.x=element_text(size=10),axis.text.y=element_text(size=15),legend.position = "none", plot.title = element_text(hjust = 0.5))
sexism_hip.graph<-sexism_hip.graph+xlab("State")+ylab("Hippocampal Volume")+ylim(6000,10000)+ggtitle("Sex/Gender Structural Stigma")
phvol.sex<-mean(sexism_hip$hipvol1)+ -29.561*unique(sexism_hip$sexism_factor)[order(unique(sexism_hip$sexism_factor))]

#Add a point for the hippocampal volume in that state as predicted by structural stigma when all covariates are mean centered.
sexism_hip.graph<-sexism_hip.graph+geom_point(aes(x=1,y=phvol.sex[1]))+geom_point(aes(x=2,y=phvol.sex[2]))+geom_point(aes(x=3,y=phvol.sex[3]))+geom_point(aes(x=4,y=phvol.sex[4]))+geom_point(aes(x=5,y=phvol.sex[5]))+geom_point(aes(x=6,y=phvol.sex[6]))+geom_point(aes(x=7,y=phvol.sex[7]))+geom_point(aes(x=8,y=phvol.sex[8]))+geom_point(aes(x=9,y=phvol.sex[9]))+geom_point(aes(x=10,y=phvol.sex[10]))+geom_point(aes(x=11,y=phvol.sex[11]))+geom_point(aes(x=12,y=phvol.sex[12]))+geom_point(aes(x=13,y=phvol.sex[13]))+geom_point(aes(x=14,y=phvol.sex[14]))+geom_point(aes(x=15,y=phvol.sex[15]))+geom_point(aes(x=16,y=phvol.sex[16]))+geom_point(aes(x=17,y=phvol.sex[17]))

#Latinx structural stigma
#Create ordered categorical variables of states in order of structural stigma
immigrant_hip$state<-as.factor(immigrant_hip$immigrant_factor)
statelev<-NULL
for(x in 1:17){
 statelev[x]<-ss_hip$STATE[which(as.factor(ss_hip$immigrant_factor)==levels(immigrant_hip$state)[x])][1]
 }
levels(immigrant_hip$state)<-levels(ss_hip$STATE)[statelev]

#Graph violin plots
immigrant_hip.graph<-ggplot(aes(x=state,y=hipvol1,fill=state),data=immigrant_hip[which(immigrant_hip$latim==0),])+geom_violin()+theme_classic()+theme(text = element_text(size=20),axis.text.x=element_text(size=10),axis.text.y=element_text(size=15),axis.title.y=element_blank(), legend.position = "none", plot.title = element_text(hjust = 0.5))
immigrant_hip.graph<-immigrant_hip.graph+xlab("State")+ylab("Hippocampal Volume")+ylim(6000,10000)+ggtitle("Latinx Structural Stigma")
phvol.lat<-mean(immigrant_hip$hipvol1)+ -41.022*unique(immigrant_hip$immigrant_factor)[order(unique(immigrant_hip$immigrant_factor))]
#Add a point for the hippocampal volume in that state as predicted by structural stigma when all covariates are mean centered.
immigrant_hip.graph<-immigrant_hip.graph+geom_point(aes(x=1,y=phvol.lat[1]))+geom_point(aes(x=2,y=phvol.lat[2]))+geom_point(aes(x=3,y=phvol.lat[3]))+geom_point(aes(x=4,y=phvol.lat[4]))+geom_point(aes(x=5,y=phvol.lat[5]))+geom_point(aes(x=6,y=phvol.lat[6]))+geom_point(aes(x=7,y=phvol.lat[7]))+geom_point(aes(x=8,y=phvol.lat[8]))+geom_point(aes(x=9,y=phvol.lat[9]))+geom_point(aes(x=10,y=phvol.lat[10]))+geom_point(aes(x=11,y=phvol.lat[11]))+geom_point(aes(x=12,y=phvol.lat[12]))+geom_point(aes(x=13,y=phvol.lat[13]))+geom_point(aes(x=14,y=phvol.lat[14]))+geom_point(aes(x=15,y=phvol.lat[15]))+geom_point(aes(x=16,y=phvol.lat[16]))+geom_point(aes(x=17,y=phvol.lat[17]))

#Race structural stigma
#Create ordered categorical variables of states in order of structural stigma
racism_hip$state<-as.factor(racism_hip$racism_factor)
statelev<-NULL
for(x in 1:17){
 statelev[x]<-ss_hip$STATE[which(as.factor(ss_hip$racism_factor)==levels(racism_hip$state)[x])][1]
 }
levels(racism_hip$state)<-levels(ss_hip$STATE)[statelev]

#Graph violin plots
racism_hip.graph<-ggplot(aes(x=state,y=hipvol1,fill=state),data=racism_hip)+geom_violin()+theme_classic()+theme(text = element_text(size=20),axis.text.x=element_text(size=10),axis.text.y=element_text(size=15),axis.title.y=element_blank(), legend.position = "none", plot.title = element_text(hjust = 0.5))
racism_hip.graph<-racism_hip.graph+xlab("State")+ylab("Hippocampal Volume")+ylim(6000,10000)+ggtitle("Race Structural Stigma")
phvol.race<-mean(racism_hip$hipvol1)+ -57.268*unique(racism_hip$racism_factor)[order(unique(racism_hip$racism_factor))]
#Add a point for the hippocampal volume in that state as predicted by structural stigma when all covariates are mean centered.
racism_hip.graph<-racism_hip.graph+geom_point(aes(x=1,y=phvol.race[1]))+geom_point(aes(x=2,y=phvol.race[2]))+geom_point(aes(x=3,y=phvol.race[3]))+geom_point(aes(x=4,y=phvol.race[4]))+geom_point(aes(x=5,y=phvol.race[5]))+geom_point(aes(x=6,y=phvol.race[6]))+geom_point(aes(x=7,y=phvol.race[7]))+geom_point(aes(x=8,y=phvol.race[8]))+geom_point(aes(x=9,y=phvol.race[9]))+geom_point(aes(x=10,y=phvol.race[10]))+geom_point(aes(x=11,y=phvol.race[11]))+geom_point(aes(x=12,y=phvol.race[12]))+geom_point(aes(x=13,y=phvol.race[13]))+geom_point(aes(x=14,y=phvol.race[14]))+geom_point(aes(x=15,y=phvol.race[15]))+geom_point(aes(x=16,y=phvol.race[16]))+geom_point(aes(x=17,y=phvol.race[17]))
